# Evolutionary analysis of the exocyst in streptophytes links EXO70 diversification to dominance over SEC3 in membrane targeting

**DOI:** 10.64898/2026.01.28.702365

**Authors:** Samuel Haluška, Edita Janková-Drdová, Matěj Drs, George Alexandru Caldarescu, Roman Skokan, Přemysl Pejchar, Viktor Žárský, Martin Potocký

## Abstract

The exocyst is a conserved octameric vesicle-tethering complex essential for targeted secretion. It is organized into two modules (I: SEC3, SEC5, SEC6, SEC8; II: SEC10, SEC15, EXO70, EXO84). In plants, the module II subunits SEC15, EXO84, and especially EXO70 have diversified into multiple subfamilies, yet the evolutionary origins and functional consequences of this diversification remain unclear. Here we reconstruct exocyst evolution across streptophytes using phylogenomic, functional complementation, and structural modeling analyses. We show that the three major EXO70 subfamilies originated in anydrophytes—the common ancestor of Zygnematophyceae and land plants—indicating that EXO70 diversification had begun at the dawn of plant terrestrialization. Complementation of Arabidopsis *exo70* mutants with EXO70 paralogs from the liverwort *Marchantia polymorpha* and the streptophyte alga *Klebsormidium nitens* demonstrates that the ancestral canonical function is retained in the EXO70.1 lineage, whereas other subfamilies have undergone substantial functional specialization. We further uncover an evolutionary shift in exocyst membrane targeting: while *Klebsormidium* SEC3 retains autonomous membrane-recruitment capacity, land-plant SEC3 subunits have lost this ability, rendering exocyst targeting increasingly dependent on EXO70. Together, these findings suggest that early EXO70 diversification, combined with the redistribution of membrane-targeting functions within the exocyst, enabled paralog-specific exocyst recruitment and facilitated the emergence of specialized secretion pathways during plant terrestrialization.

**Significance Statement:** The exocyst is a conserved protein complex that targets secretory vesicles to the plasma membrane across eukaryotes. In land plants, the EXO70 subunit diversified extensively, but the origins and consequences of this diversification remained unclear. We show that three major EXO70 subfamilies arose in the common ancestor of land plants and their closest algal relatives, at the dawn of plant terrestrialization. Cross-species complementation reveals that the EXO70.1 lineage retains the ancestral canonical function, whereas other subfamilies specialized. We further uncover an evolutionary shift in the mechanism of exocyst membrane targeting, from SEC3 toward EXO70, that could have enabled distinct EXO70 paralogs to direct secretion to different cellular sites during the colonization of land.

## Introduction

The exocyst complex is an evolutionarily conserved, hetero-octameric tethering complex that directs secretory vesicles to the plasma membrane (PM) prior to SNARE-mediated membrane fusion (1–3). Composed of eight subunits—SEC3, SEC5, SEC6, SEC8, SEC10, SEC15, EXO70, and EXO84—the exocyst is organized into two structural and functional modules: module I (SEC3, SEC5, SEC6, SEC8) and module II (SEC10, SEC15, EXO70, EXO84), which assemble via N-terminal CorEX helical bundles (4–6). While this modular exocyst architecture, dating to the last eukaryotic common ancestor, is preserved across diverse lineages, the contributions of individual subunits to complex function, their evolutionary diversification, and their functional conservation in plants remain largely unresolved.

In plants, the exocyst is essential for polarized growth, cell morphogenesis, and cytokinesis (7–11), and has also been implicated in autophagy-related processes (12, 13). In the model angiosperm *Arabidopsis thaliana*, multiple exocyst subunits accumulate at sites of intense secretion, including the forming cell plate during cytokinesis (14). Exocyst subunits also co-localize as discrete foci at exocytically active plasma membrane domains in growing root epidermal cells (9). Genetic disruption of core exocyst subunits, including SEC3, SEC6, SEC8, SEC15, or EXO84, results in severe developmental defects, underscoring the conserved requirement for an intact exocyst complex in plant growth and viability (15, 16).

While exocyst subunits are encoded by a single gene in yeast or by a few paralogs in metazoans, the land plant exocyst exhibits a striking feature: expansion of the EXO70 family to 23 paralogs in Arabidopsis, or 47 paralogs in rice (17, 18). Previous phylogenetic analyses have identified three major land-plant EXO70 subfamilies, proposed to have emerged around the divergence of streptophyte algae and land plant (embryophyte) lineages (18, 19). In contrast, other angiosperm exocyst subunits (except EXO84) are encoded by three or fewer paralogs (18). Among Arabidopsis EXO70 paralogs, the EXO70A clade (belonging to the EXO70.1 subfamily) has been the most extensively studied due to its near-ubiquitous expression pattern and the severe phenotypic deviations of *exo70A1* loss-of-function mutants. These include dwarfism, loss of apical dominance, disruption of polarized cell expansion in root hairs, and near-complete sterility (15, 16). Another EXO70 isoform, EXO70B1, is implicated in autophagy: loss-of-function mutants show reduced vacuole anthocyanin accumulation and spontaneous hypersensitive response-like lesions on leaves (12). Isoform EXO70H4 is involved in Arabidopsis trichome secondary cell wall biogenesis by delivering callose to a specific plasma membrane—cell wall domain (20, 21).

A key mechanistic feature of the exocyst is that its membrane targeting—the process by which the complex is recruited to specific PM domains—strongly depends on interactions between specific subunits and negatively charged phospholipids. In yeast and mammals, both EXO70 and SEC3 serve as membrane-targeting landmarks through their phosphoinositide-binding surfaces (22–26). These interactions position the exocyst at sites of polarized secretion, enabling small GTPase-regulated vesicle delivery and localized cell expansion (1, 27, 28). Importantly, the electrostatic properties of protein surfaces determine the affinity and specificity of membrane association. Because membrane-binding surfaces are also often involved in protein–protein interactions within the exocyst core, electrostatic properties represent an evolutionarily tunable parameter that can simultaneously influence membrane targeting and inter-subunit interfaces (29, 30).

In angiosperms, the balance between EXO70 and SEC3 in membrane targeting differs markedly from the yeast paradigm. Although Arabidopsis SEC3 retains a lipid-binding PH domain, its contribution to exocyst PM localization is minor (31); instead, EXO70A1 dominates exocyst membrane recruitment in the Arabidopsis sporophyte (6). Similarly, in rice, the localization and function of the OsSEC3a subunit in crown roots depend on lipid-driven PM recruitment of OsEXO70A1 (32). Crucially, the evolutionary timing, functional consequences, and mechanistic underpinnings of the shift in landmarking roles between SEC3 and EXO70 in plants remains unknown.

Over the past decade, considerable effort has been put into understanding the evolution of exocyst complex proteins in land plants, particularly the highly diversified EXO70 subunit (19, 33, 34). With the increasing availability of recent genomic and transcriptomic data from various streptophyte lineages (35–39), we sought to reassess the plant exocyst evolution and diversification with improved resolution.

Here, we combined phylogenetic reconstruction, cross-species complementation assays, and structural bioinformatics approaches to examine the evolution and diversification of the exocyst complex across streptophytes. By focusing on SEC3, SEC15 and EXO70 homologs from three species spanning the Streptophyta lineage—the streptophyte alga *Klebsormidium nitens*, the liverwort *Marchantia polymorpha*, and the angiosperm *Arabidopsis thaliana*—we analyzed the conservation and divergence patterns within the exocyst and experimentally tested functional relationships underlying plasma membrane targeting. Our results indicate that during plant terrestrialization, subunits from EXO70.1 subfamily became the dominant determinant of exocyst plasma membrane localization at the expense of SEC3.

## Results

### Exocyst evolution in plants is marked by extensive expansion of module II subunits in streptophytes

To trace exocyst evolution across streptophytes, we collected the exocyst complex subunits homologs from all currently sequenced streptophyte algae and selected embryophytes (Table S1) and reconstructed their phylogenetic relationships (Fig. 1A, Fig. S1). Analysis of the two modules that constitute plant exocyst complex revealed that the genes coding the subunits of the module II expanded extensively during plant evolution. Specifically, three of four subunits—SEC15, EXO70 and EXO84—underwent paralog diversification at distinct stages of streptophyte evolution (Fig. S1). EXO84 subunit at first diversified into two subfamilies – EXO84AB and EXO84CXC – in the common ancestor of tracheophytes, followed by later duplications and losses (Fig. S1I). SEC15 diverged into two paralogous lineages – SEC15A and SEC15B – in the ancestor of spermatophytes (Fig. S1F), consistent with the previous reports (40). Most notably, EXO70 – the most expanded exocyst subunit – underwent repeated multiplication and fixation events beginning already in anydrophytes (the common ancestor of *Zygnematophyceae* and Embryophyta). Importantly, these expansion events continued in all currently most species-rich plant lineages (Fig. 1A, Fig. S1G-H): in addition to flowering plants, significant EXO70 expansion has occurred also in mosses (bryophytes), and even in desmids (zygnematophytes).

**Figure 1.**
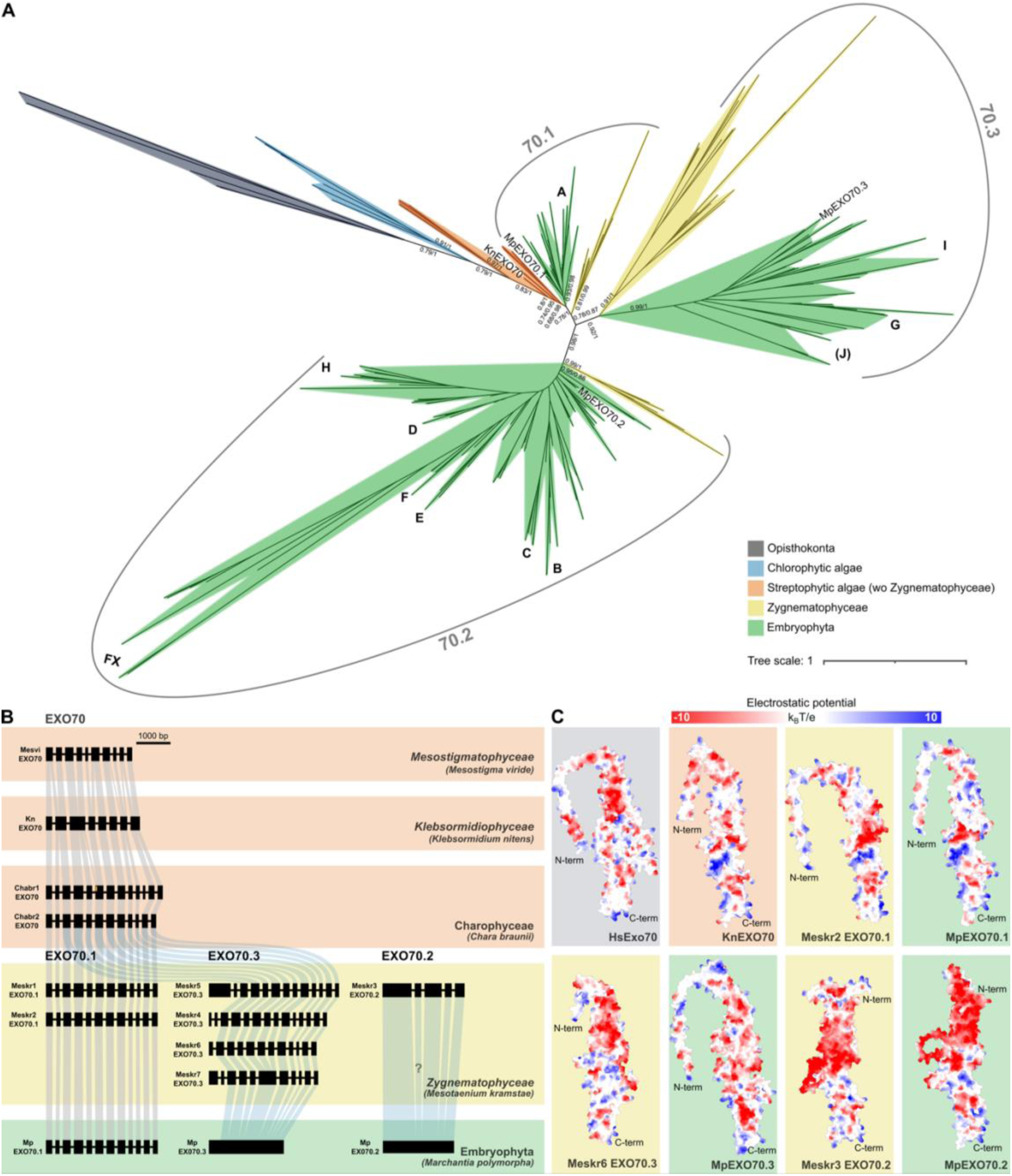
Deep origin and functional diversification of EXO70 subfamilies in streptophytes. (**A**) Phylogenetic analysis of EXO70 proteins across Viridiplantae and representative opisthokont outgroups (see also Fig. S1 for details). (**B**) Analysis of exon–intron architecture in EXO70 homologs from representative streptophyte lineages. For clarity, intron lengths are not drawn to scale; Fig. S2 shows the true scale. (**C**) Molecular surface models of selected EXO70 proteins illustrating electrostatic potential distributions across evolutionary lineages and subfamilies. Surface charge is shown from negative (red) to positive (blue) electrostatic potential. Chabr, *Chara braunii*; Hs, *Homo sapiens*; Kn, *Klebsormidium nitens*; Mp, *Marchantia polymorpha*; Meskr, *Mesotaenium kramstae*; Mesvi, *Mesostigma viride*.

Collectively, our analyses reveal a strikingly uneven evolutionary history of the plant exocyst, in which diversification is concentrated primarily in module II, whose subunits expanded across streptophyte evolution. These findings identify module II as the principal evolutionary substrate for functional innovation in the plant exocyst complex.

### EXO70 triplication predates embryophytes with independent expansions in several Anydrophyta lineages

With the increasing availability of genomes and transcriptomes from these previously undersampled groups (Table S1), we were able to reexamine the EXO70 diversification events in more detail. Strikingly, we found that EXO70 homologs from conjugating algae (*Zygnematophyceae*) cluster within the three well-defined EXO70 subfamilies originally described in embryophytes (Fig. 1A). This finding extends the origin of three EXO70 subfamilies by tens of millions of years (41) to the anydrophyte lineage (42). Accordingly, the number of EXO70 paralogs in this group can be found from 7 in *Mesotaenium kramstae* (Fig. 1; Fig. S1, S2) to possibly 68 in the *Penium margaritaceum* genome (43). Notably, an independent EXO70 duplication occurred also within the *Charophyceae*—two Chara EXO70 homologs are shared also with their *Nitella* relative (Fig. S1). Additionally, our data strongly put the root of the EXO70 expansion next to the EXO70.1 family, as summarily indicated by the phylogenetic position orthologs from non-plant models, green algae and pre-anydrophyta streptophyte algae (Fig. 1A, Supplementary Dataset. S1).

To gain sequence-independent insight into the early EXO70 evolution, we also analyzed the conservation of exon-intron structure of EXO70s encoding genes from selected members of streptophyte lineage. These data corroborate the evolutionary position of EXO70.1 subfamily (including *Zygnematophyceae*) close to other streptophyte algae and also indicate contrasting modes of evolution between the three subfamilies. While EXO70.1 and EXO70.3 subfamilies evolve primarily by gene duplication, the most diversified subfamily—EXO70.2—likely proliferated through reverse transcription followed by genomic integration as most representatives of the EXO70.2 subfamily are intronless genes (Fig. 1B, Fig. S2).

Following the initial triplication in anydrophyta, EXO70 copy number remains low in several early-diverging embryophyte lineages. Hornworts, liverworts, and lycophytes typically encode 3–5 EXO70 homologs (one or two per subfamily), whereas extensive expansion in crown mosses is largely associated with whole-genome duplication events (33). Major expansion of the EXO70 family begins in Euphyllophytes. The stepwise manner of these events is evident in Fig. 1 and Fig. S1, which also reveals the loss of one evolutionary conserved EXO70.3 subfamily member (that we termed clade J) in Angiosperms.

Notably, the diversification of EXO70 subunits sequences results in many instances in large shifts in electrostatic charge distribution along the rod-like body of EXO70 molecules. This tendency is most prominent among the 70.2 subfamily representatives (Fig.1) where N-terminal part of the protein features prominent negative charge, largely deviating also from the non-plant EXO70s (Fig. 1C).

Taken together, our data demonstrate that EXO70 diversification began early in streptophyte evolution, with the establishment of three major subfamilies happening prior to embryophytes, followed by lineage-specific expansions, distinct duplication mechanisms, and pronounced sequence and structural divergence.

### Marchantia encodes three EXO70 genes with distinct localization and overexpression phenotypes

The liverwort *Marchantia polymorpha* possesses only three EXO70 homologs, one representing each of the three EXO70 subfamilies. We took advantage of this minimal EXO70 complement, together with the emerging status of *Marchantia* as a model plant (44), to investigate the evolutionary conservation of EXO70 localization and function using GFP fusion proteins and overexpression approaches. All three *Marchantia* EXO70 genes were N-terminally fused to GFP and expressed under the control of the strong endogenous MpEF1α promoter (45).

The evolutionarily conserved MpEXO70.1 isoform (homologous to AtEXO70A isoforms; Fig. 1) localized exclusively to the plasma membrane in thalli, rhizoid initials and rhizoids. In contrast, MpEXO70.2 exhibited predominantly cytoplasmic localization, while MpEXO70.3 displayed a dual plasma membrane–cytoplasmic localization pattern (Fig. 2A, B; Fig. S3A, B). Overexpression of MpEXO70.1 and MpEXO70.3 did not produce obvious alterations in overall thallus morphology. By contrast, overexpression of MpEXO70.2 caused severe disruption of thallus architecture, including tearing of the upper epidermis and strongly impaired chloroplast biogenesis. Regenerated thalli of MpEXO70.2 overexpressors were abnormally twisted and lacked gemma cups, with gemmae instead developing directly on the top of the thallus (Fig. 2C; compare also with Fig. S3C).

**Figure 2.**
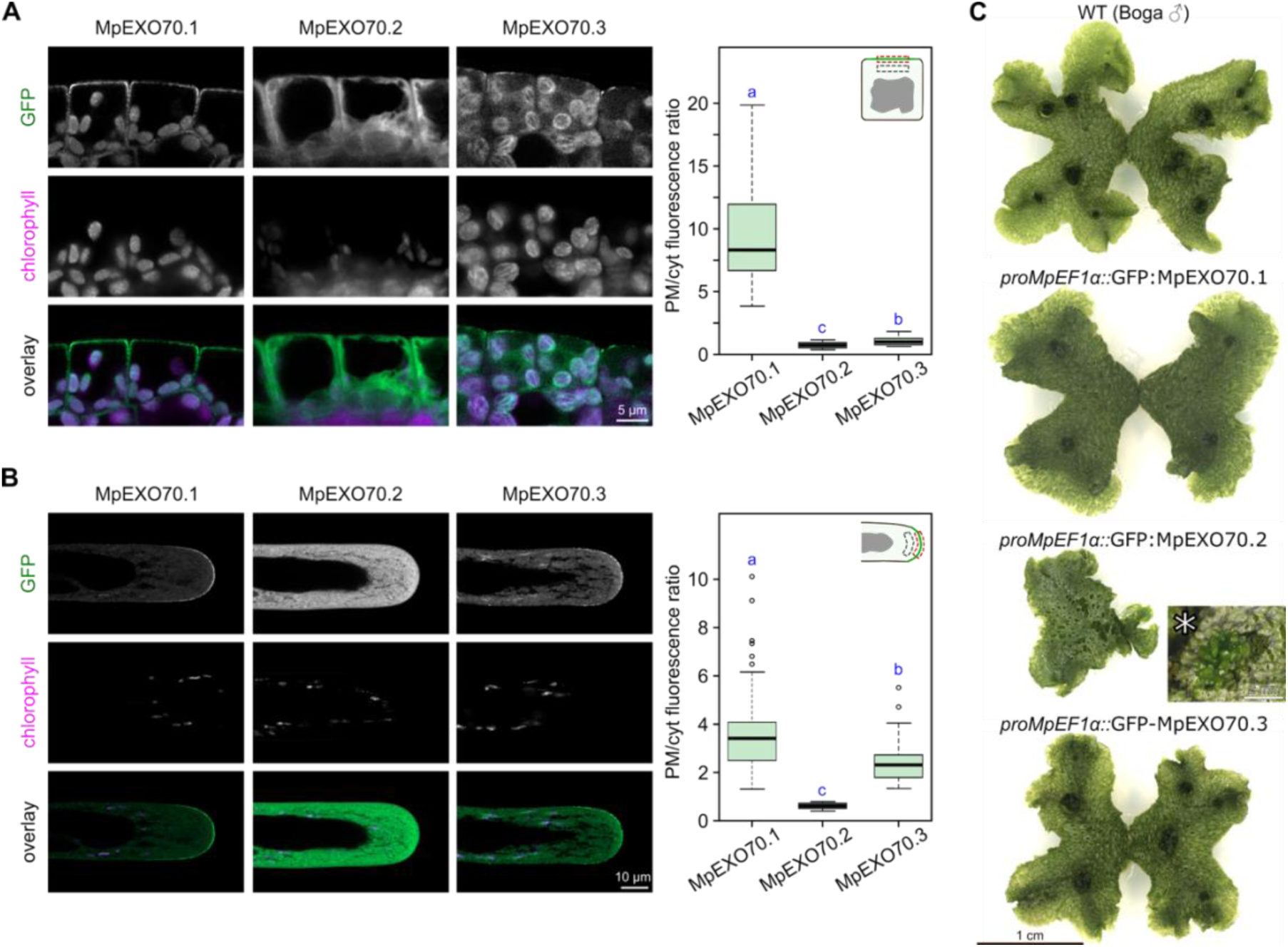
Localization and overexpression analyses of Marchantia EXO70 subfamily representatives in *M. polymorpha*. Localization of GFP-tagged MpEXO isoforms in (**A**) gemmae, (**B**) rhizoids. GFP, chlorophyll autofluorescence and overlay channels are shown to highlight the true GFP signal. Panels on the right represent the quantitative analysis of the PM to cytoplasm (PM/Cyt) fluorescence ratio at the indicated regions. Distinct letters denote statistically different groups (Kruskal-Wallis test with Holm correction, p < 0.05). For each line, at least 50 cells from ca. 20 independent gemmae from two independent lines were quantified. (**C**) 3 weeks old *in vitro* Marchantia thallus morphology carrying over-expression constructs. Asterisk denotes a detailed view of delayed gemmae emergence in EXO70.2 over-expressor after 8 weeks of cultivation. See also Fig. S3 for additional details.

Collectively, our observations highlight pronounced functional and localization differences between the evolutionarily conserved EXO70.1 and the more divergent EXO70.2 and EXO70.3 isoforms in *M. polymorpha*, that are consistent with the overall evolutionary pattern of the three major EXO70 subfamilies.

### SEC15 subunit shows deep functional conservation across streptophytes

Because SEC15 is markedly more conserved than the highly diversified EXO70 family, we used it as a model subunit to test functional conservation across streptophytes. Across 39 streptophyte sequences, SEC15 proteins share 66% amino acid identity and 79% similarity, and have a mean protein length of 808 ± 43 amino acids (Supplementary Dataset S2). This conservation, together with the clear sporophytic phenotypes of Arabidopsis *sec15b* mutants (40), made SEC15 a suitable model subunit for testing functional conservation across streptophytes.

We therefore examined SEC15 orthologs from *A. thaliana*, *M. polymorpha*, and *K. nitens*, representing angiosperms, bryophytes (as their most distant land plant sister group), and streptophyte algae, respectively. When expressed under the control of the AtUBQ10 promoter in Arabidopsis, both the Marchantia and Klebsormidium SEC15 homologs localized prominently to the lateral plasma membrane domains of rhizodermal cells and to developing cell plates, closely mirroring the localization pattern of AtSEC15b (Fig. 3A). Importantly, both orthologs fully complemented the short etiolated hypocotyl phenotype of the Arabidopsis *sec15b* mutant and restored the overall growth defects associated with loss of AtSEC15b function (Fig. 3B, C).

**Figure 3.**
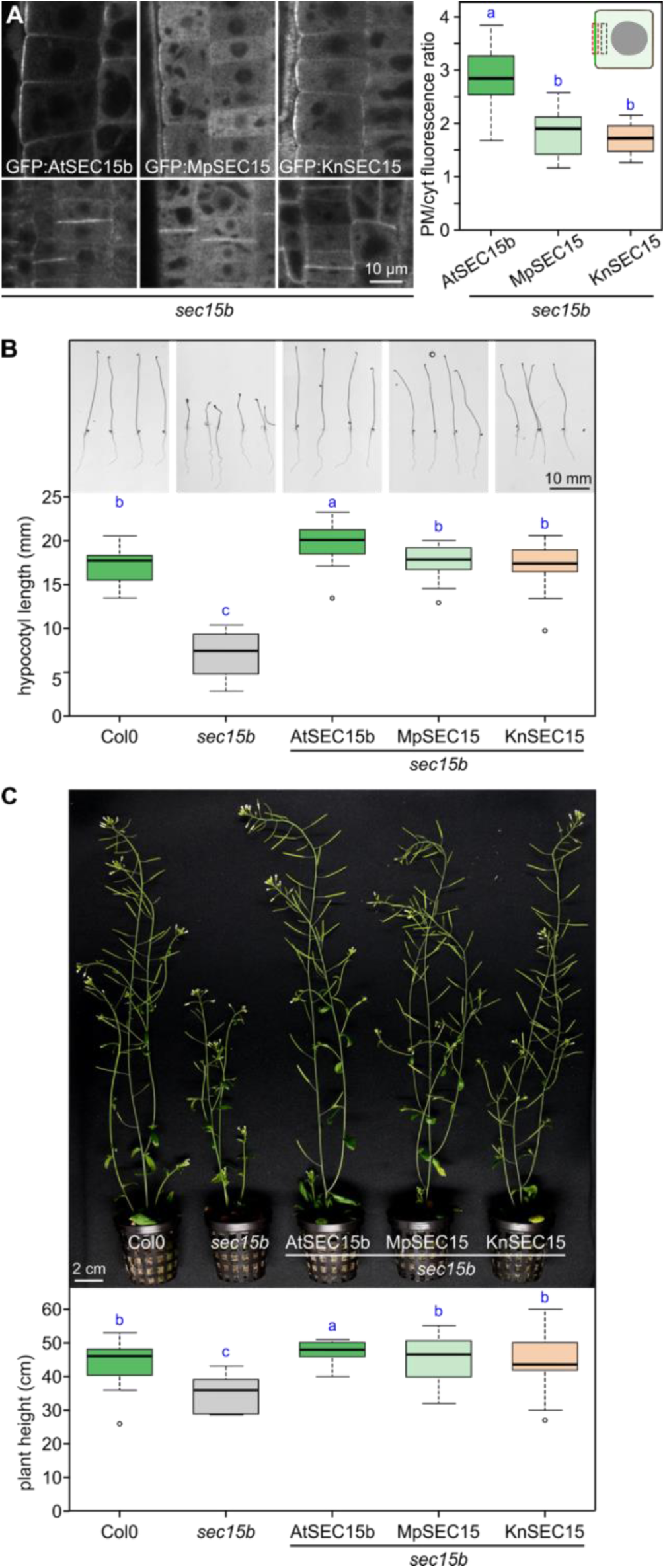
SEC15 orthologs across streptophytes display deep localization and functional conservation. (**A**) Localization of GFP-tagged SEC15 isoforms from Arabidopsis, Marchantia and Klebsormidium in root epidermal cells, focusing on lateral plasma membrane domain (upper left panels) and cell plate (bottom left panels). The constructs were expressed in the *sec15b* mutant background under the control of AtUBQ10 promoter. Panel on the right represents the quantitative analysis of the PM to cytoplasm (PM/Cyt) fluorescence ratio at the indicated region. For each line, at least 12 cells from four independent plants were quantified. (**B**) Representative images and length of 7-day-old etiolated hypocotyls of WT, *sec15b* or *sec15b* seedlings expressing GFP-tagged AtSEC15b, MpSEC15 or KnSEC15. For each line, 20 seedlings were quantified. (**C**) Representative images and heights of 6-week-old WT, *sec15b* or *sec15b* plants expressing GFP-tagged AtSEC15b, MpSEC15 or KnSEC15. For each line, at least 9 plants were quantified. In all plots, distinct letters denote statistically different groups (Kruskal-Wallis test with Holm correction, p < 0.05).

Together, these results demonstrate that molecular conservation of the SEC15 exocyst subunit correlates strongly with functional conservation across the streptophyte lineage, validating cross-complementation as a robust approach for evolutionary conservation studies.

### EXO70 diversification reveals ancestral conservation and lineage-specific functional specialization in streptophytes

Having established SEC15 as a deeply conserved exocyst subunit whose molecular conservation directly translates into functional interchangeability across streptophytes, we next asked how this paradigm applies to EXO70—the most diversified exocyst component in plants. Since EXO70 underwent early gene multiplication into three distinct subfamilies, the question arises: how does this evolutionary expansion relate to functional conservation and specialization, both within and across EXO70 subfamilies?

To address this, we first examined the subcellular localization and complementation capacity of all three Marchantia polymorpha EXO70 paralogs in Arabidopsis wild-type plants and the *exo70A1* mutant background. All three Marchantia genes were cloned under the constitutive AtUBQ10 promoter, fused to GFP at the N-terminus, and expressed in Arabidopsis. Subcellular localization was subsequently analyzed in root epidermal cells. Consistent with observations in Marchantia, MpEXO70.1 displayed strong enrichment at the lateral plasma membrane domain of rhizodermal cells (Fig. 4A, Fig. S4A). Importantly, MpEXO70.1 complemented the severe developmental defects of the Arabidopsis *exo70A1* mutant, restoring overall adult plant architecture and normal developmental progression (Fig. 4B; Fig. S4B). Moreover, MpEXO70.1 rescued the impaired hypocotyl elongation in dark-grown seedlings and restored normal root hair growth (Fig. 4C, D).

**Figure 4.**
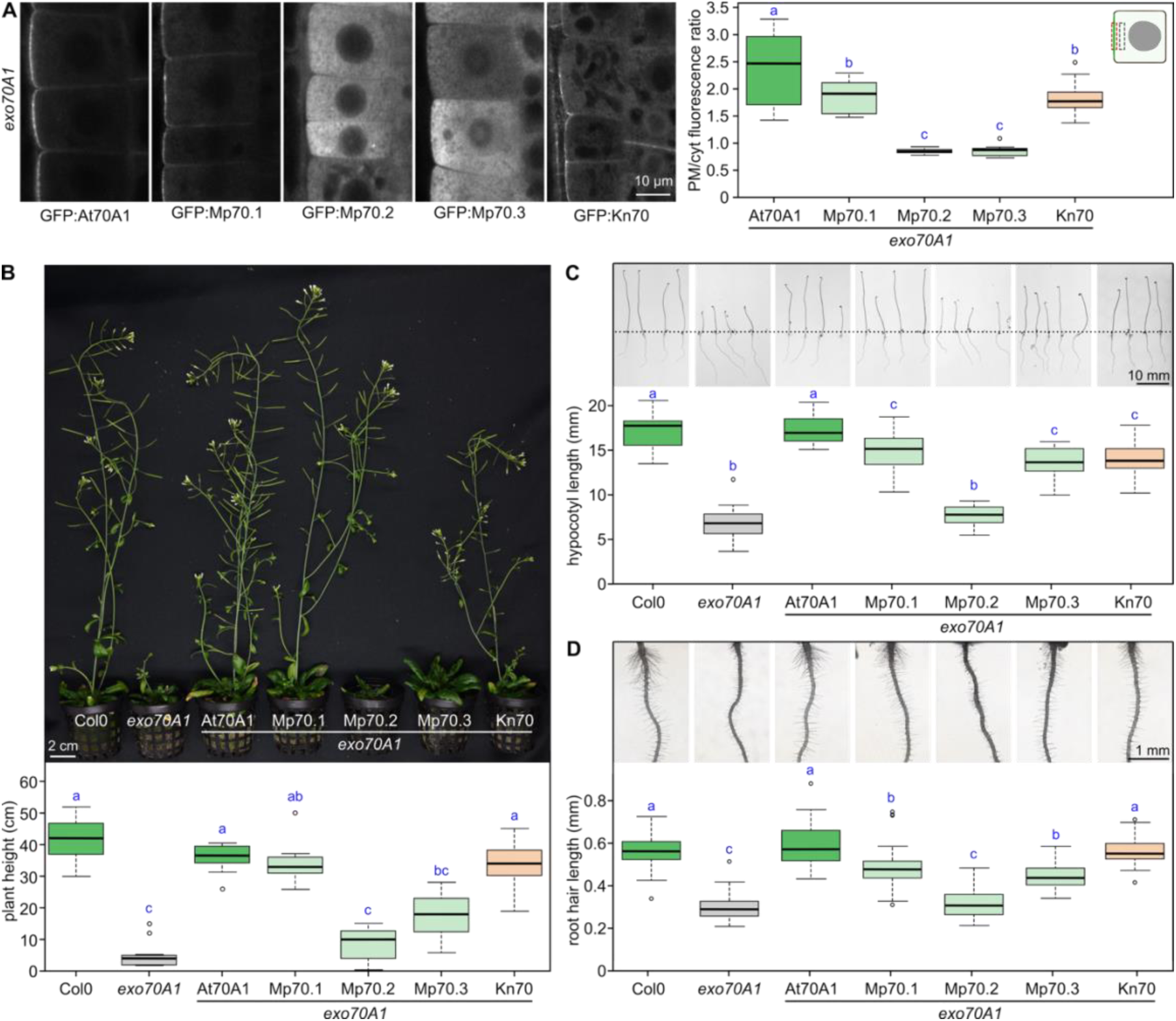
Localization and complementation analyses of Marchantia and Klebsormidium EXO70 homologs in Arabidopsis. **(A**) localization of GFP-tagged Arabidopsis EXO70A1, all three EXO70 orthologues from *M. polymorpha* and a single EXO70 ortholog from *K. nitens* in Arabidopsis roots of *exo70A1* mutant line. Analysis of the PM to cytoplasm (PM/Cyt) fluorescence ratio at the outer lateral PM in *exo70A1* root epidermal cells. For each line, at least 12 cells from four independent plants were quantified. (**B**) Expression of GFP-tagged MpEXO70.1 and KnEXO70 largely rescues the dwarf and fertility phenotype of *exo70A1* mutant. For each line, at least 9 plants were quantified. (**C**) Multiple GFP-tagged Marchantia EXO70 and KnEXO70 restore the hypocotyl elongation defect of the *exo70A1* mutant. For each line, 20 seedlings were quantified. (**D**) Marchantia and Klebsormidium GFP-tagged EXO70 orthologs display distinct rescue effects on the short root hair growth defect characteristic for the *exo70A1* mutant. For each line, the 5 longest roothairs from 10 seedlings were quantified. In all plots, distinct letters denote statistically different groups (Kruskal-Wallis test with Holm correction, p < 0.05). See Fig. 4 for additional details.

Interestingly, MpEXO70.3 expression resulted in partial complementation across all assessed traits. Although GFP:MpEXO70.3 localized mostly to cytoplasm and MpEXO70.3-expressing plants remained sterile, exhibited delayed inflorescence initiation, and showed compromised apical dominance similar to the *exo70A1* mutant, they attained greater final height than the mutant, albeit with greater variability than MpEXO70.1-expressing plants (Fig. 4B; Fig. S4B). Partial rescue was also observed for hypocotyl elongation in etiolated seedlings and for root hair development (Fig. 4C, D). In sharp contrast, MpEXO70.2, which also localized predominantly to the cytoplasm, failed to complement any of the major *exo70A1* mutant phenotypes (Fig. 4B–D; Fig. S4B).

We next investigated the evolutionary depth of functional conservation within the EXO70.1 subfamily. Phylogenetic and structural homology analyses indicated that single EXO70 algal homologues predating the EXO70 triplication cluster close to the EXO70.1 subfamily (Fig. 1A), suggesting that this subfamily may retain the single ancestral canonical exocyst subunit function. To test this hypothesis experimentally, we cloned the sole EXO70 homolog from the filamentous streptophyte alga *K. nitens*. When expressed in Arabidopsis under the AtUBQ10 promoter, KnEXO70 localized to both the cytoplasm and characteristic exocyst-associated plasma membrane domains at the lateral surface of root epidermal cells, as well as to developing cell plates during cytokinesis (Fig. 4A). Strikingly, Klebsormidium EXO70 complemented all major developmental defects of the Arabidopsis *exo70A1* mutant, including impaired overall growth, hypocotyl elongation, and root hair formation (Fig. 4B–D; Fig. S4B).

One defining feature controlling EXO70 functionality is its localized electrostatic charge, which is crucial for membrane binding and interactions with other exocyst subunits (6, 30, 46). We therefore sought to analyze charge distribution across streptophyte EXO70 paralogs and test whether it serves as a distinguishing factor in EXO70 diversification. To this end, we generated AlphaFold models for 59 distinct EXO70 proteins and quantitatively compared their electrostatic surface properties using Protein Interaction Property Similarity Analysis (PIPSA) (47), thereby assessing charge distribution similarities and differences across paralogs. PIPSA clustering clearly distinguished members of the three EXO70 subfamilies. Strikingly, single-copy EXO70 paralogs from pre-anydrophyta streptophyte algae (including Klebsormidium) clustered together with land plant EXO70.1 members, consistent with our complementation analysis (Fig. S5). Surprisingly, EXO70.2 and EXO70.3 members from Amborella, Marchantia, and *Zygnematophyceae* also clustered together with their respective Arabidopsis orthologs, despite their greater evolutionary distance (Fig. S1, S5).

We therefore next tested whether functional conservation of Marchantia EXO70 paralogs extends beyond the EXO70.1 subfamily and whether the single Klebsormidium EXO70 can function in multiple subfamily contexts. To this end, we assessed their capacity to complement phenotypes of two well-characterized Arabidopsis mutants from the EXO70.2 subfamily: *exo70H4*, which fails to deposit secondary cell wall thickenings in trichomes (21), and *exo70B1*, which manifests through early senescence and an anthocyanin accumulation defect (12). However, none of the tested Marchantia or Klebsormidium EXO70 paralogs could complement the *exo70H4* or *exo70B1* mutant phenotypes (Fig. S4C–E), indicating further clade-specific functional diversification within the EXO70.2 subfamily.

Together, these results demonstrate that EXO70 diversification in land plants led to pronounced functional divergence among paralogs. The canonical EXO70 function—underpinned by conserved surface charge distribution—was retained by the EXO70.1 lineage and is already present in streptophyte algae, predating plant terrestrialization. Conversely, the specialized functions of Arabidopsis EXO70.2 subfamily members could not be complemented by either MpEXO70.2 or KnEXO70, indicating that these functions evolved independently in distinct land plant lineages.

### SEC3 retains ancestral membrane targeting capacity in streptophytes but becomes EXO70-dependent in embryophytes

Studies in fungal and animal systems have established the exocyst subunit SEC3, together with EXO70, as a key determinant of exocyst targeting to specific plasma membrane domains. In these systems, membrane targeting relies on coordinated interactions with membrane lipids and small GTPases, providing a critical “landmarking” function for exocyst assembly and activity (23–25). In contrast, although SEC3 remains essential for exocyst function in plants, its landmarking role appears diminished in Arabidopsis (31). Building on our analyses of SEC15 and EXO70, we therefore sought to comparatively assess the functional contribution of SEC3 to exocyst localization and activity across the streptophyte lineage. To this end, we transformed heterozygous Arabidopsis *sec3a* mutants with GFP-tagged SEC3 homologs from *A. thaliana*, *M. polymorpha*, and *K. nitens*, expressed under the constitutive AtUBQ10 promoter.

SEC3 homologs from all three species exhibited highly similar localization patterns, accumulating at the lateral plasma membrane domains of root epidermal cells, at maturing cell plates, and at the tips of growing pollen tubes (Fig. 5A). Consistent with this conserved subcellular localization, both Marchantia and Klebsormidium SEC3 homologs fully complemented the Arabidopsis *sec3a* pollen transmission defect and restored wild-type growth (Fig. 5B; Fig. S6), demonstrating deep functional conservation of SEC3 across streptophytes.

**Figure 5.**
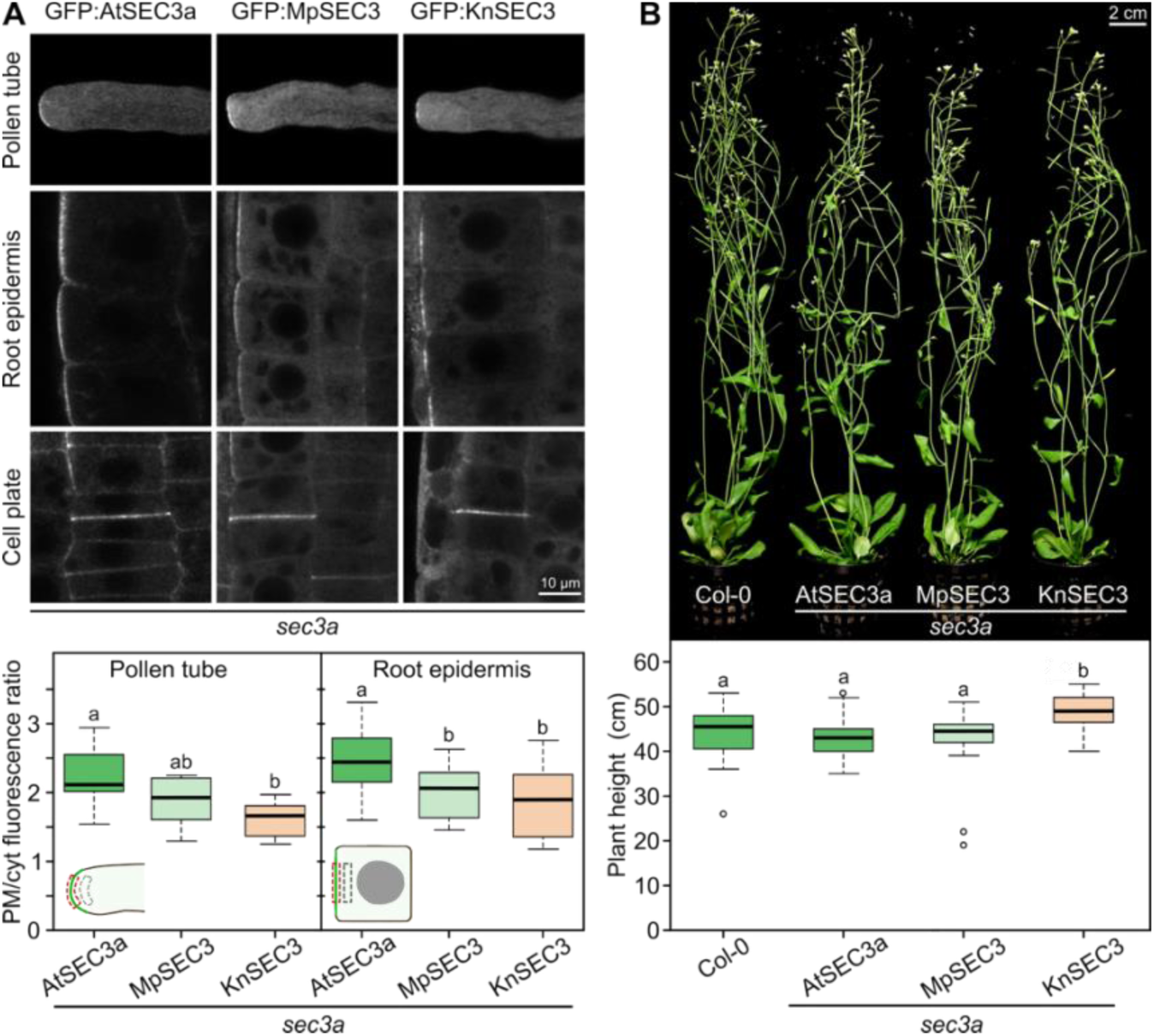
Localization and complementation analyses of native, Marchantia and Klebsormidium SEC3 homologs in Arabidopsis *sec3a* mutant. (**A**) Localization of GFP-tagged SEC3 isoforms from Arabidopsis, Marchantia and Klebsormidium in growing pollen tubes, lateral plasma membrane domain of root epidermal cells, and cell plates. The constructs were expressed in the *sec3a* mutant background under the control of AtUBQ10 promoter. The panel below represents the analysis of the PM to cytoplasm (PM/Cyt) fluorescence ratio in pollen tubes and root epidermis at indicated regions. For each line, at least 6 pollen tubes were quantified and 12 root epidermal cells from four independent plants were quantified. (**B**) Representative images and heights of 6-week-old WT, and *sec3a* plants expressing GFP-tagged AtSEC3a, MpSEC3 or KnSEC3. For each line, at least 15 plants were quantified. In all plots, distinct letters denote statistically different groups (Kruskal-Wallis test with Holm correction, p < 0.05). See Fig. S6 for more details.

To assess how the relative contributions of SEC3 and EXO70 to exocyst plasma membrane targeting evolved, we next examined the behavior of SEC3 homologs in the Arabidopsis *exo70A1* mutant background, extending previous observations indicating an increased reliance on EXO70A1 for plasma membrane targeting in land plants (6, 31). Strikingly, whereas the Arabidopsis and Marchantia SEC3 proteins were displaced from the plasma membrane and accumulated in cytoplasmic patches in the *exo70A1* mutant (identical to those previously described in (6)), Klebsormidium SEC3 retained partial plasma membrane localization in the absence of EXO70A1 (Fig. 6A, B). We confirmed that this residual membrane association occurred within the context of the assembled exocyst complex, as evidenced by co-immunoprecipitation of endogenous SEC6 with Klebsormidium SEC3 (Fig. 6C).

**Figure 6.**
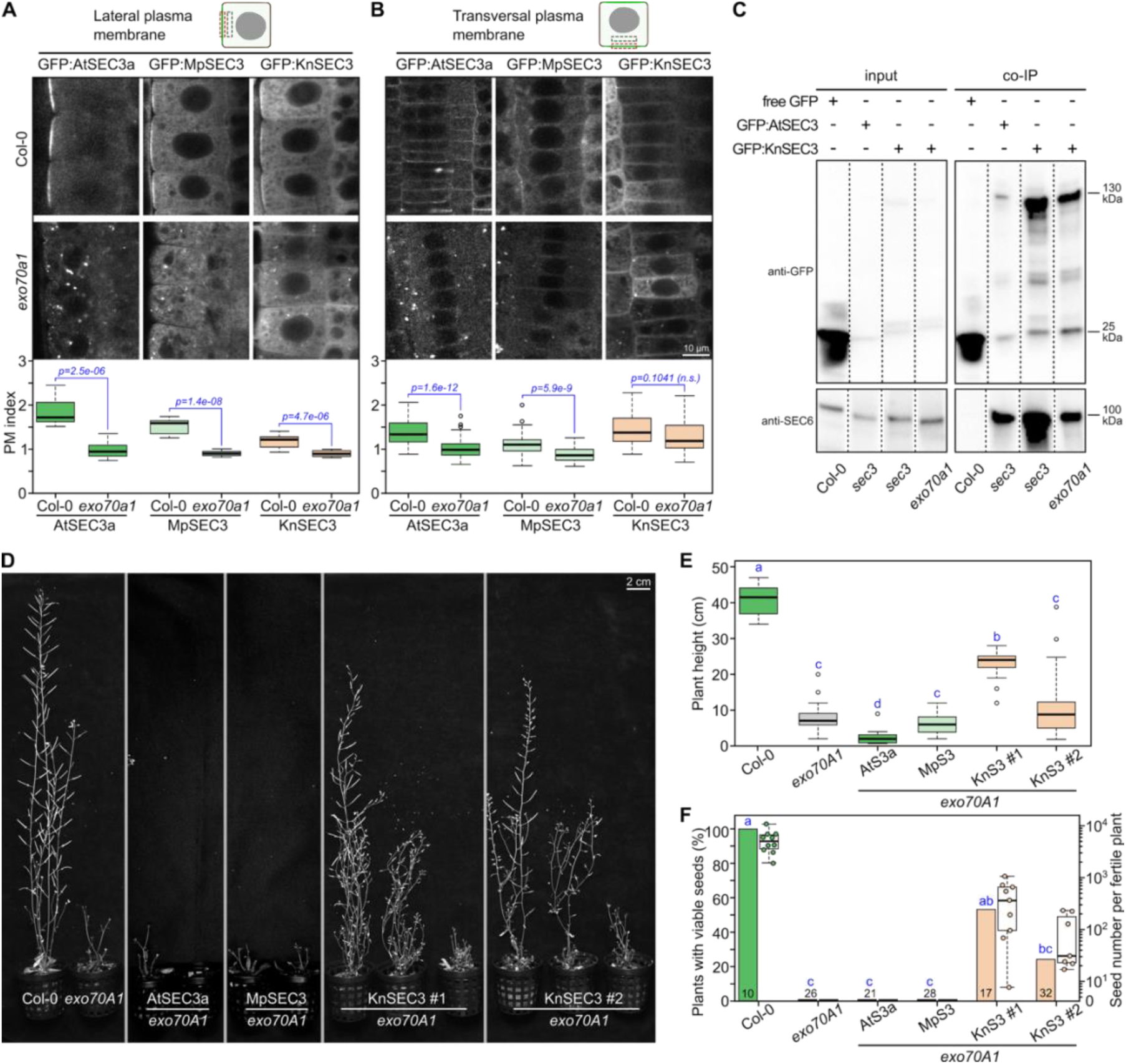
Evolutionary shift in SEC3-mediated exocyst membrane targeting. (**A, B**) Localization of GFP-tagged SEC3 isoforms from Arabidopsis, Marchantia, and Klebsormidium in wild-type and *exo70A1* mutant backgrounds. PM-to-cytoplasm (PM/Cyt) fluorescence ratio was analyzed at the outer lateral PM in root epidermal cells (**A**) and the transversal PM in cortical cells (**B**). Klebsormidium SEC3 still partially localizes to the plasma membrane in Arabidopsis *exo70A1*. Statistical significance from Mann-Whitney U tests is indicated. (**C**) Co-immunoprecipitation of the exocyst core subunit SEC6 with GFP-tagged AtSEC3a and KnSEC3 in wild-type, *sec3a*, and *exo70A1* mutant backgrounds. Detection of SEC6 in SEC3 co-immunoprecipitates confirms association of both Arabidopsis and Klebsormidium SEC3 with the exocyst complex, including in the *exo70A1* background. Input and co-IP controls are shown. (**D, E, F**) Klebsormidium SEC3 partially complements *exo70A1* growth and fertility defects. Representative plant images (**D**) and quantitative analysis of plant height (**E**) and fertility (**F**) show significant rescue by two independent KnSEC3 lines compared with *exo70A1* mutants and with Arabidopsis or Marchantia SEC3 expression. The number of plants analyzed is indicated at the bottom of the graph. Distinct letters above boxplots denote statistically different groups (Kruskal-Wallis test with Holm correction, p < 0.05). See Fig. S6 for additional details.

Consistent with these localization differences, independent Klebsormidium SEC3 lines partially but robustly complemented the severe growth and developmental defects of the Arabidopsis *exo70A1* mutant, including restoration of fertility and seed production. In contrast, neither Arabidopsis nor Marchantia SEC3 homologs provided detectable complementation in the *exo70A1* background (Fig. 6D–F).

Taken together, these results indicate that while SEC3 remains functionally compatible with exocyst complex across streptophytes, its autonomous contribution to exocyst plasma membrane targeting has been reduced during land plant evolution. We propose that, concomitant with the multiplication and diversification of EXO70 paralogs, EXO70.1 became the dominant determinant of exocyst membrane targeting in embryophytes, whereas SEC3 retained this capability only in streptophyte lineages from before EXO70 multiplication events.

### Comparative structural modeling reveals how Klebsormidium SEC3 preserves exocyst function in the absence of EXO70

To understand why Klebsormidium SEC3 rescues the Arabidopsis *exo70a1* mutant phenotypes more effectively than land-plant SEC3 orthologs, we first examined its sequence and structural features. Multiple-sequence alignment did not identify motifs that distinguish Klebsormidium SEC3 from land-plant SEC3 proteins. Likewise, surface analysis of the N-terminal PH domain, which is important for membrane binding in yeast but dispensable in Arabidopsis (25, 31), did not suggest altered membrane-binding properties of the Klebsormidium protein (Fig. S7). Together, these results argued against a simple sequence- or surface-based explanation and suggested that the beneficial effect of Klebsormidium SEC3 emerges in the context of the assembled exocyst complex. We therefore used AlphaFold3 (48), followed by comparative structural analysis of predicted exocyst assemblies.

Using a workflow benchmarked on the yeast exocyst structure (Fig. S8), we predicted near full-length exocyst complexes from Arabidopsis, Marchantia, and Klebsormidium, as well as Arabidopsis complexes lacking EXO70A1 and carrying SEC3 from Arabidopsis, Marchantia, or Klebsormidium. All predictions converged on a similar overall architecture comprising two modules connected primarily via interfaces involving the well-supported subunits SEC5, SEC8, SEC15, and EXO84. In contrast, SEC3 (module I) and EXO70 (module II)—and, to a lesser extent, SEC6 (module I) and SEC10 (module II)—, appeared tightly linked through their N-terminal CorEX motifs but were otherwise positionally less supported, indicating their structural flexibility. Predictions containing EXO70 subfamily 70.2 and 70.3 members suggested even greater positional variability, especially for EXO70.2 proteins (Fig. S8).

We next compared the Arabidopsis-derived assemblies lacking EXO70A1. These models remained globally similar to the native Arabidopsis complex, but they showed higher ipSAE (interaction prediction Score from Aligned Errors) (49) subunit-subunit scores, consistent with a more compact arrangement and reduced separation between the two modules (Fig. S9A–D). To interpret these structural changes, we defined two functional complexes—the native Arabidopsis exocyst and the EXO70A1-deficient complex carrying Klebsormidium SEC3—and two non-functional complexes—EXO70A1-deficient assemblies containing either Arabidopsis SEC3 or Marchantia SEC3—based on the experimentally observed rescue phenotypes and performed their comparative analysis.

This comparison revealed that the non-functional complexes formed novel SEC5–SEC10 and SEC5–EXO84 interfaces that were absent from both functional variants (Fig. 7E). These aberrant contacts were concentrated in the N-terminal CorEX motifs and led to stronger coupling between module I and module II. By contrast, the corresponding interfaces were much looser in the functional complexes (Fig. 7H). A similarly low number of inter-module contacts was also observed in the native Klebsormidium exocyst model (Fig. 7G; Fig. S9E). Although we could not pinpoint a single sequence feature responsible for this behavior, we noted that the N-terminal region of Klebsormidium SEC3 near the affected SEC5 region is enriched in small, uncharged residues relative to the corresponding regions of land-plant SEC3 orthologs, potentially favoring a less constrained packing at this interface.

**Figure 7.**
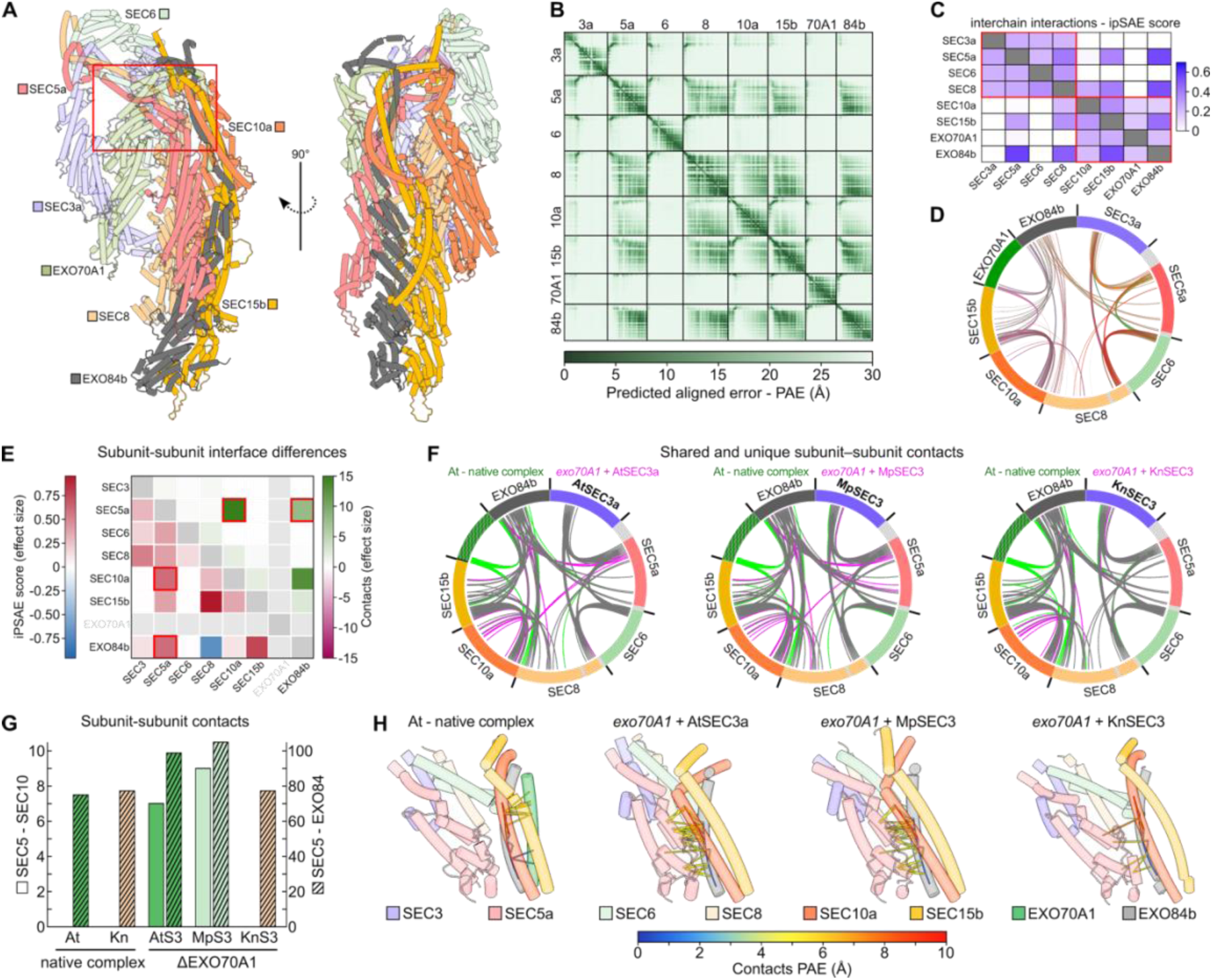
Comparative structural analysis of exocyst complexes reveals a unique role for Klebsormidium SEC3. **(A)** Schematic representations of the AlphaFold3 (AF3) exocyst model from *A. thaliana*. Subunits with uncertain global positions are shown as semi-transparent. The red rectangle indicates the position of the CorEX helical bundles. **(B)** Predicted aligned error (PAE) plots for the Arabidopsis exocyst from the AF3 prediction, showing confidence in relative residue positioning. Low (dark green) and high (light green) PAE values indicate robustly predicted rigid regions and flexible or uncertain regions, respectively. **(C)** Heatmap of ipSAE (interaction prediction Score from Aligned Errors) values for individual subunit–subunit interfaces in the Arabidopsis exocyst, computed with a 10 Å threshold. Red rectangles indicate interactions within modules I and II. **(D)** Chord plot projection of inter-subunit contacts in the Arabidopsis exocyst complex. Inter-subunit residue–residue contacts with distance < 5 Å and predicted aligned error (PAE) < 10 Å are shown. Regions trimmed before modelling are shown in grey. **(E)** Heatmap of differences in ipSAE metrics and inter-subunit contacts between functional and non-functional exocyst variants, presented as Cohen’s *d* effect sizes. Red rectangles indicate SEC5–SEC10 and SEC5–EXO84 interactions. **(F)** Differential inter-subunit contacts between models of the native Arabidopsis complex and variants lacking EXO70A1 and containing SEC3 from Arabidopsis, *Marchantia*, or *Klebsormidium*. Shared contacts are grey, contacts specific to the native complex are green, and contacts specific to the ΔEXO70A1 variants are magenta. **(G)** Number of interface contacts between SEC5–SEC10 and SEC5–EXO84 subunits. **(H)** Close-up of robustly predicted contacts between subunits of modules I and II. The region shown corresponds to the boxed area in panel A. Only contacts with distance < 5 Å and PAE < 10 Å are shown.

Together, our comparative structural analysis suggests that the interface between exocyst modules I and II—mediated by CorEX motifs of SEC5, SEC10, and EXO84—is an important determinant of exocyst functionality in the absence of EXO70. In the chimeric, EXO70-deficient exocyst, Klebsormidium SEC3 prevents the formation of aberrant, overly stabilizing inter-module contacts, thereby maintaining a looser architecture compatible with native exocyst function. These findings suggest that in land-plant exocyst, the structural interplay between SEC3 and EXO70 has been tuned to regulate inter-module dynamics, and that the unique structural properties of Klebsormidium SEC3—likely reflecting the ancestral state prior to the expansion of the EXO70 family—can bypass the requirement for EXO70-mediated regulation of this interface.

## Discussion

### Exocyst evolution in streptophytes reveals asymmetric constraint within a conserved complex

The exocyst is a conserved tethering complex whose core subunit composition predates the divergence of major eukaryotic lineages (34), while retaining fundamental roles in exocytosis and possibly also in autophagy (13, 50). Our streptophyte-wide analysis reveals, however, that individual exocyst subunits have experienced markedly different evolutionary constraints. These differences converge at the level of the two exocyst modules and define two principal evolutionary axes discussed below: diversification of EXO70 within module II, and a plant-specific redistribution of membrane-targeting roles between SEC3 and EXO70.

Across all streptophyte lineages, the copy number of module I subunits mostly remains constrained to one or two per subunit, whereas most module II subunits diversified extensively during plant evolution (Fig. 8A). The earliest and most pronounced expansion involves EXO70, beginning in anydrophytes, followed by duplication of EXO84 at the base of tracheophytes and subsequent diversification in the most species-rich groups of plants (Fig. 8A; Fig. S1G-I). A further module II innovation is the duplication of SEC15 in spermatophytes. While the functional significance of EXO84 duplication remains unclear, we hypothesize that SEC15 duplication in seed plants may be linked to increasing functional differentiation between sporophytic and gametophytic programs and to processes associated with sexual reproduction (40). Together, this asymmetry indicates that plant exocyst evolution proceeded not by uniform modification of the entire complex, but by the generation of multiple module II variants that interact with a comparatively constrained module I to support distinct cellular functions.

**Figure 8.**
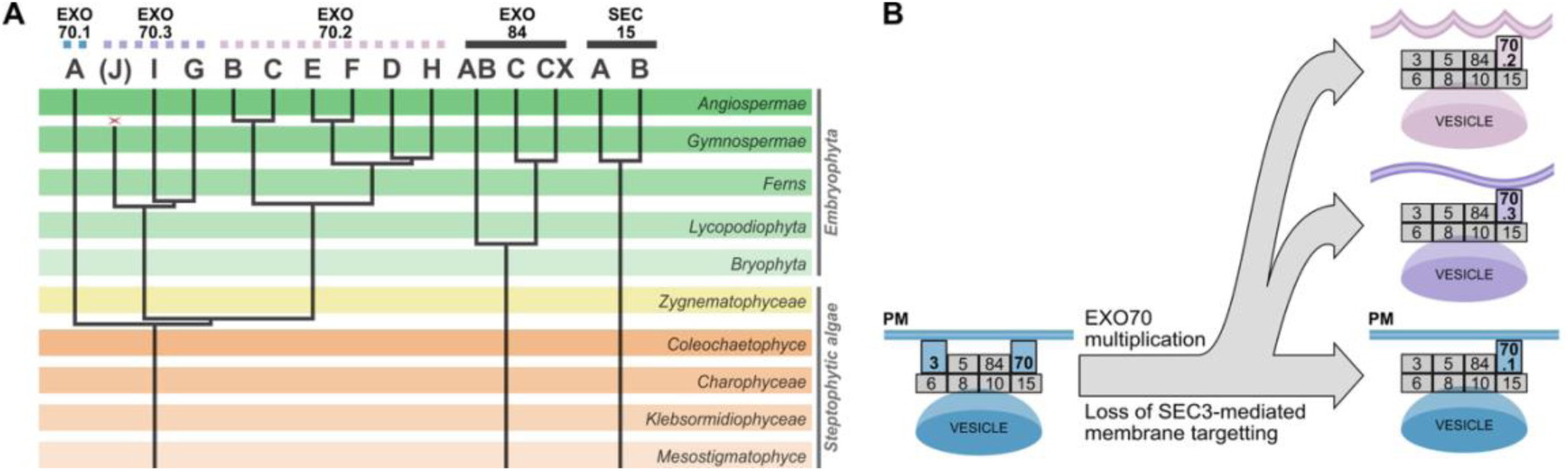
Early EXO70 diversification shapes plant exocyst function. **(A)** Schematic summary of cladistic phylogenetic analyses illustrating the emergence and diversification of selected module II exocyst subunits (EXO70, EXO84, and SEC15) during streptophyte evolution. (**B**) Hypothetical model depicting a plant-specific evolutionary shift of exocyst membrane-targeting mechanisms. In evolutionary distant streptophyte algae, SEC3 contributes directly to plasma membrane (PM) association of the exocyst complex, together with EXO70. During embryophyte evolution, this contribution is reduced, coincident with EXO70 multiplication and functional specialization. In land plants, EXO70.1 emerges as the dominant determinant of exocyst membrane localization to plasma membrane, while SEC3 becomes increasingly dependent on EXO70 for effective PM targeting. This redistribution is proposed to have enabled paralog-specific recruitment of the exocyst complex or exocyst-derived subassemblies to distinct membrane domains.

### Early EXO70 multiplication predates embryophytes and canonical function retains in EXO70.1 subfamily

Previous phylogenetic studies placed the emergence of the three major EXO70 subfamilies in the common ancestor of embryophytes (15, 18, 19). Our analyses extend this origin deeper, to anydrophytes—the common ancestor of embryophytes and zygnematophytes (42). This timing places EXO70 diversification at a critical evolutionary transition at the dawn of land colonization. In anydrophytes, duplication of an ancestral single EXO70 was rapidly followed by a second duplication, yielding the three subfamilies—EXO70.1, EXO70.2, and EXO70.3—whose topology is conserved in both zygnematophytes and embryophytes (Fig. 1, 8A).

To test functional consequences of this diversification, we exploited cross-species complementation using SEC3, SEC15, and EXO70—subunits representing contrasting evolutionary trajectories from both modules—and three phylogenetically distant species: *K. nitens* (streptophyte alga), *M. polymorpha* (bryophyte), and *A. thaliana* (tracheophyte). Single homologs of SEC3 and SEC15 from Klebsormidium and Marchantia localized to the plasma membrane in Arabidopsis and fully complemented *sec3a* and *sec15b* mutants, confirming deep functional conservation of these subunits. In contrast, EXO70 paralogs showed striking functional divergence. Klebsormidium EXO70 and Marchantia EXO70.1 complemented the *exo70A1* mutant, whereas MpEXO70.3 showed partial localization and rescue, and cytoplasmic MpEXO70.2 failed to complement.

These results demonstrate that canonical EXO70 function in the whole complex was already established in streptophyte algae and retained in the EXO70.1 lineage. Notably, although algal EXO70 proteins fall outside the EXO70.1 subfamily phylogenetically, they cluster with EXO70.1 in protein surface charge analyses, underscoring the importance of conserved electrostatics for membrane association and protein–protein interactions (29). Together with correlating localization patterns and contrasting overexpression phenotypes of the three paralogs in Marchantia, these findings reveal deep functional divergence among the basal EXO70 subfamilies.

### EXO70 diversification and exocyst association

Recently, De la Concepcion et al. (30) proposed that EXO70 diversification was enabled by electrostatic shifts in the CorEX N-terminal region that reduce its stable association with the exocyst (“complex escape”), with evolutionarily later reassociation invoked to explain exocyst binding by some Arabidopsis EXO70.2 members. In Marchantia, this model is supported by the predominantly cytosolic localization of EXO70.2 during cytokinesis, contrasted with EXO70.1 and EXO70.3 enrichment at the cell plate alongside core subunits.

Our data are consistent with several core observations of this framework, including the functional and localization separation of the three Marchantia paralogs and the superior ability of MpEXO70.1 (and partially MpEXO70.3) to substitute for canonical EXO70A1-dependent functions in Arabidopsis. At the same time, evidence from angiosperms indicates that the relationship between EXO70 diversification and exocyst association is more heterogeneous than a simple dichotomy between complex-bound and complex-free states. Several EXO70.2 paralogs implicated in defense, autophagy, and specialized secretion (EXO70B1, EXO70B2, EXO70E2) show physical or functional association with core subunits, whereas others (EXO70C1/C2) appear largely excluded from the complex and may antagonize secretion (12, 51–55). This pattern points to fine-grained specialization in the degree and context of exocyst engagement within the EXO70.2 subfamily.

Importantly, exocyst-independent EXO70 functions do not necessarily require permanent dissociation from the complex. In opisthokonts, EXO70 exists in both free and exocyst-incorporated pools, with the free pool contributing to membrane deformation and curvature generation (56, 57). EXO70 paralogs may diversify primarily by modulating their equilibrium between free and complex-associated states, rather than by complete evolutionary disengagement.

Plant exocyst function also extends beyond canonical secretion associated with primary cell wall formation. Functional specificity among EXO70.2 paralogs is particularly striking, as closely related isoforms are frequently non-interchangeable in vivo, indicating specialization for distinct trafficking pathways and membrane domains (21, 58, 59). We hypothesize that this specialization reflects differential recruitment of exocyst assemblies to pathway-specific membrane environments rather than uniform “escape” from the complex.

Collectively, these findings support a model in which early EXO70 triplication provided a platform for functional diversification of exocyst activity. Differential electrostatic properties of EXO70 paralogs likely modulate the stability, geometry, and context of exocyst association, enabling lineage- and pathway-specific solutions. Within this view, the complex escape model and our data represent complementary perspectives on how electrostatic tuning and regulatory plasticity shaped EXO70 evolution in plants.

### Plant-specific redistribution of exocyst membrane targeting: from SEC3 contribution to EXO70.1 dominance

In parallel with EXO70 multiplication, we identify a second major evolutionary transition: a weakening of SEC3’s autonomous contribution to plasma membrane targeting in favor of EXO70.1 dominance. This redistribution emerges as a defining feature of embryophyte exocyst organization.

Early evidence for this shift came from studies showing that the N-terminal PH domain of Arabidopsis SEC3a, previously assumed to mediate membrane binding, is dispensable for SEC3 function. An N-terminally truncated SEC3 lacking membrane-binding capacity fully complemented *sec3a* pollen tube defects (31). Consistently, plasma membrane localization of SEC3a—and of the exocyst complex as a whole—is strictly dependent on EXO70.1 in Arabidopsis (6), a dependency we now extend to Marchantia. A direct interaction between SEC3 and EXO70A1, first reported in Arabidopsis (7) and absent from opisthokont systems, further supports a plant-specific strengthening of SEC3–EXO70 coupling. Recent work in rice independently corroborates this conclusion by showing that SEC3a function in crown root formation depends on EXO70A1 association (32).

Strikingly, this dependence does not apply to the evolutionarily distant streptophyte alga Klebsormidium. In an Arabidopsis *exo70A1* background, Klebsormidium SEC3 retains partial plasma membrane localization and partially rescues mutant phenotypes. Comparative AF3-based modeling suggests a mechanistic explanation: while both exocyst modules form stable cores across species, SEC3 and EXO70 occupy flexible global positions. The presence of Klebsormidium SEC3 results in fewer stabilizing inter-module contacts, leading to a looser architecture that may permit partial membrane landmarking in the absence of EXO70A1. Although predictive, these observations suggest that SEC3 sequential or structural features can tune inter-module plasticity and thereby modulate reliance on EXO70.

In summary, we conclude that embryophyte exocyst evolution involved a coordinated process in which EXO70 triplication coincided with a redistribution of membrane-targeting capacity from SEC3 to EXO70.1 (Fig. 8B). This shift likely enabled different EXO70 paralogs to direct the exocyst complex—or exocyst-derived subassemblies—to distinct membrane domains and cellular processes.

## Materials and Methods

### Sequence retrieval and analyses

The goal was to acquire at least two species from each major evolutionary group (namely *Mesostigmatophyceae*, *Klebsormidiophyceae*, *Charophyceae*, *Zygnematophyceae*, *Anthocerotophyta*, *Marchantiophyta*, true *Bryophyta*, *Lycopodiophyta*, *Monilophyta* (Ferns), *Gymnosperms* and *Angiosperms*), to create balanced and largely representative datasets. The sequences have been retrieved from publicly available genome and transcriptome databases (Table S1). Sequences from the genome releases were revisited with transcriptome assemblies within the identical species or genus and occasionally accordingly reassembled using FGENESH+ (60). MUSCLE (61) or Mafft (62) algorithms were used for the multiple alignments generation, and maximum likelihood algorithm in PhyML (63) was employed for the phylogenetic tree reconstruction with LG+G4 settings, implemented in the SeaView multiplatform program (64).

### Constructions of binary vectors

To fit our needs, pMpGWB101 binary vector (Addgene #68555) has been cut by *BamHI* and *SacI* and modified by Annealed Oligo Cloning (using P1+P2, Table S2.). Subsequently, *proMpEF1α* (P3+P4, Table S2.), *proAtUBQ10* (P5+6, Table S2.) and GFP (P7+P8, Table S2.) have been PCR amplified and ligated into the backbone using *HindIII*, *NcoI* and *BamHI* restriction enzymes. All the exocyst subunits were ligated N-terminally to the GFP using *BamHI/XmaI*-CDS-*PacI* restriction sites (P11-P36, Table S2).

### Plant materials and growth conditions

The *M. polymorpha* BoGa male strain has been maintained on ½-strength Gamborg medium (65) solidified with 1.2% agar (Duchefa Biochemie) in a cultivation room at 22 °C under a 16/8 h light/dark regime with a light intensity of approximately 185 µmol photons m⁻² s⁻¹. For the transformation, G – AgarTrap method has been used (66) with selection on 10 μg.ml^−1^ Hygromycin and 100 μg.ml^−1^ Claforan.

*A. thaliana* seeds were surface sterilised, stratified for 2d at 4°C and germinated on vertical ½ Murashige-Skoog supplemented with 1% sucrose, vitamin mixture, and 1.6% plant agar (Duchefa Biochemie) at the same regime as mentioned above. Young seedlings were transferred to pellets (Jiffy Products International) and moved to *ex vitro* (22 °C, 16/8 light/dark regime). In this study Arabidopsis mutants *exo70A1* (SALK_014826) (15), *exo70H4* (SALK_023593) (20), *exo70B1* (GABI_156G02) (12), *sec15b* (SALK_130663) (40), and *sec3a* (GABI_652H12) (31) were used. Genotyping of mutants was performed using the following primer combinations (Table S2): *exo70A1* P31+P33, P31+P32; *exo70H4* P34+P35, P35+P33; *exo70B1* P36+P37, P37+P40; *sec15b* P38+P39, P39+P33; *sec3a* P31+P32+P36. Transgenic lines of *A. thaliana* were generated using *Agrobacterium tumefaciens* GV3101 strain by floral dip method (67). To evaluate the complementation capacity of evolutionary distinct variants of exocyst subunits, we analyzed their functionality in the second generation of transformed *exo70A1* or *sec3a* heterozygotes. GFP-positive seedlings were visually selected and transferred to soil. The phenotype was documented 8 weeks after germination. Plant height was measured at the post-flowering stage because *exo70A1* mutant plants exhibit delayed flowering.

### Arabidopsis pollen tube germination

Pollen grains were germinated on solid medium containing 10% sucrose, 0.01% H₃BO₃, 5 mM CaCl₂, 1 mM MgSO₄, 5 mM KCl, and 1.5% low-melting agarose (pH 7.5). Petri dishes with pollen grains were incubated inverted at room temperature for 3–5 h. Agarose blocks with germinated pollen tubes were excised and transferred to a microscopy chamber for observation.

### Microscopic observations and quantification

Subcellular localizations of selected fluorescently tagged exocyst subunits were examined in young Arabidopsis roots (7d) and Marchantia gemmae using ZEISS LSM 900 (JENA/Germany) equipped with Airyscan 2 using a 488/561 nm laser (objective: LD LCI Plan-Apo 40x/1,2 Imm Corr DIC M27, Plan-Apochromat 63x/1.40 Oil DIC M27, alpha Plan-Apochromat 100x/1.46 Oil DIC M27).

*In vitro* thallus of *M. polymorpha* and etiolated seedlings of *A. thaliana* were imaged on Stereomicroscope Leica M205FA (objective Plan-Apochromat 2x, camera Leica DMC6200 colour 2.2Mp).

Root hairs were imaged on Olympus BX51 stereo binocular microscope (objective Olympus UplanApo 4x NA 0,16, camera Olympus DP74).

For post-acquisition image processing and for all quantifications Fiji software was used (68). Figures were prepared in Inkscape. The PM association was calculated as a ratio of the PM/cytoplasm GFP signal based on mean gray values in narrow regions of the PM and next to it in the cytoplasm, avoiding vacuoles or nuclei.

### Co-immunoprecipitation and western blot analysis

*In vivo* interactions within the exocyst subunits were detected by co-immunoprecipitation using the µMACS GFP-Tagged Protein Isolation Kit (Miltenyi Biotec) according to the manufacturer’s instructions with only slight modification as described below. 0,6 g 10 d old *Arabidopsis* seedlings for each genotype were ground in liquid nitrogen and 1ml of Sec6/8 buffer was added (20 mM HEPES, pH 6.8, 150 mM NaCl, 1 mM EDTA, 1 mM DTT, 0.5% Tween-20, 100x protease inhibitor cocktail (Sigma-Aldrich). A volume of 100 μL of Anti-GFP magnetic beads was used and Wash Buffer 1 was replaced by Sec6/8 Buffer. Bound proteins were eluted with 100 μL of preheated elution buffer, separated by SDS–PAGE using 4–15% Mini-PROTEAN precast gels (Bio-Rad) and transferred to the membrane using the Trans-Blot Turbo Transfer System (Bio-Rad, Hercules, CA, USA). Western blot analysis was performed according to standard procedures, using the primary antibodies anti-Sec6 rabbit (oligopeptide generated antibody from Agrisera, dilution 1:10000), anti-GFP mouse (AS152987, Agrisera, dilution 1:1000) and Horseradish peroxidase-conjugated secondary antibodies (anti-rabbit 1:10000 and anti-mouse 1:10000; Promega). After the washing step the membrane was incubated with 1000 µL Amersham ECL Prime Western Blotting Detection Reagent (GE Healthcare, RPN2236) for 2 min and visualized using the ChemiDoc XRS+ Imaging system (Bio-Rad).

### *In silico* structural analyses

Structural models of near full-length exocyst complexes from *A. thaliana*, *M. polymorpha*, *K. nitens* and *Saccharomyces cerevisiae* were predicted using a local installation of the AlphaFold3 server (48). Prior to prediction, intrinsically disordered regions were removed from selected subunit sequences as follows: *Arabidopsis* SEC5a (residues 1–217 and 1043–1090) and SEC8 (1–34 and 226–307); *Marchantia* SEC5 (1–215 and 1042–1178) and SEC8 (1–8, 215–315, 397–440, and 539–610); *Klebsormidium* SEC5 (1–330, 1132–1155, and 1200–1346), SEC8 (207–310 and 536– 636), SEC10 (1–106), SEC15 (1–32 and 767–825), and EXO84 (1–30); and *Saccharomyces* SEC3 (1–68 and 248–610), SEC5 (1–70), SEC8 (1–33 and 1011–1065), SEC10 (1–58 and 482–556), and SEC15 (1–61 and 767–910). The accuracy of structural predictions and the propensity for inter-chain interactions were assessed using the interaction prediction Score from Aligned Errors (ipSAE) metric (49). Structure visualization and residue contact analyses were performed using ChimeraX (69).

## Acknowledgments

This work was supported by Charles University project GAUK 1544120 to S.H., by the Czech Science Foundation (GAČR) projects 22-35916S (to M.P.) and 23-05564S (to V.Ž.), and by the Ministry of Education, Youth and Sports (MEYS) project TowArds Next GENeration Crops, reg. no. CZ.02.01.01/00/22_008/0004581 of the ERDF Programme Johannes Amos Comenius. The Imaging Facility of the Institute of Experimental Botany CAS is supported by the MEYS CR (LM2023050 Czech-BioImaging), the Czech Academy of Sciences, and Institute of Experimental Botany CAS. Computational resources were provided by the e-INFRA CZ project (ID:90254), supported by the Ministry of Education, Youth and Sports of the Czech Republic. We thank Hana Soukupová, Lucie Brejšková and Jana Kaněrová for the technical assistance and Prof. Sabine Zachgo for the kind permission to use the *M. polymorpha* BoGa strain. Gratitude also goes to Prof. Takayuki Kohchi for making the pMpGWB101 plasmid available.

