## Supplementary Figure S1-S9 and Supplementary Table S1-S2 for "Evolutionary analysis of the exocyst in streptophytes links EXO70 diversification to dominance over SEC3 in membrane targeting"

*Martin Potocký

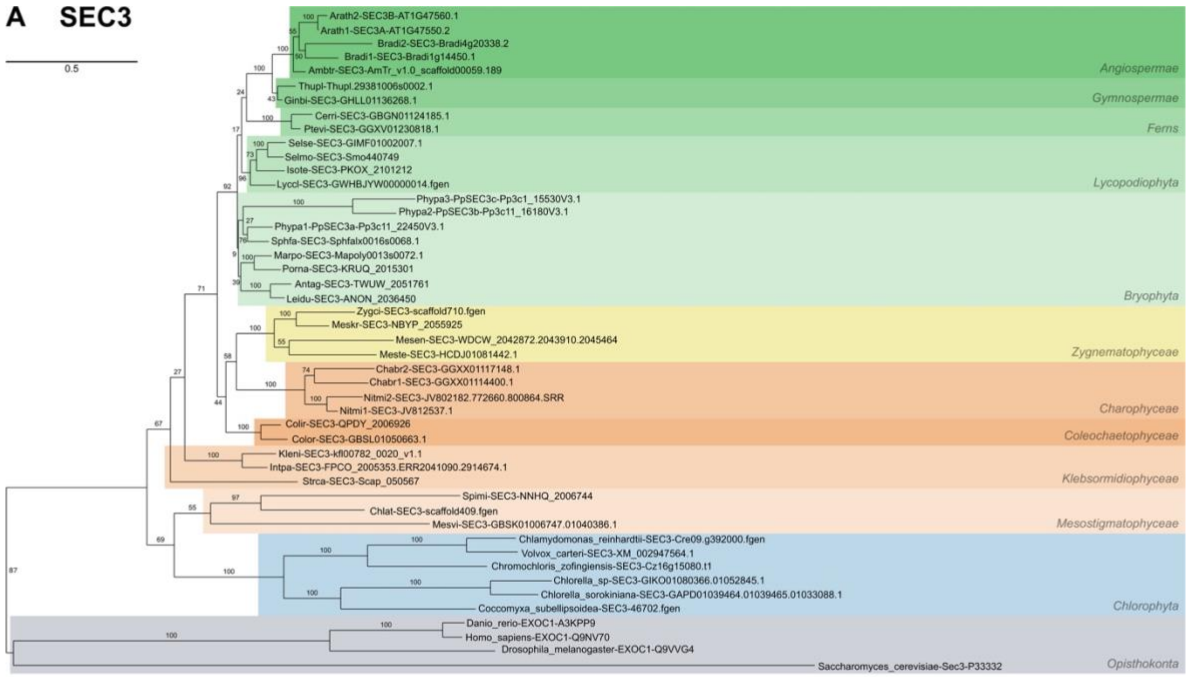
A SEC3
0.5
Arath2-SEC3B-AT1G47560.1
Arath1-SEC3A-AT1G47550.2
Bradi2-SEC3-Bradi4g20338.2
Bradi1-SEC3-Bradi1g14450.1
Ambtr-SEC3-AmTr_v1.0_scaffold00059.189
Angiospermae
Thupl-Thupl.29381006s0002.1
Ginbi-SEC3-GHLL01136268.1
Gymnospermae
Cerri-SEC3-GBGN01124185.1
Ptevi-SEC3-GGXV01230818.1
Ferns
Selse-SEC3-GIMF01002007.1
Selmo-SEC3-Smo440749
Isote-SEC3-PKOX_2101212
Lyccl-SEC3-GWHBJYW00000014.fgen
Lycopodiophyta
Phypa3-PpSEC3c-Pp3c1_15530V3.1
Phypa2-PpSEC3b-Pp3c11_16180V3.1
Phypa1-PpSEC3a-Pp3c11_22450V3.1
Sphfa-SEC3-Sphfalx0016s0068.1
Marpo-SEC3-Mapoly0013s0072.1
Porna-SEC3-KRUQ_2015301
Antag-SEC3-TWUW_2051761
Leidu-SEC3-ANON_2036450
Bryophyta
Zygci-SEC3-scaffold710.fgen
Meskr-SEC3-NBYP_2055925
Mesen-SEC3-WDCW_2042872.2043910.2045464
Meste-SEC3-HCDJ01081442.1
Zygnematophyceae
Chabr2-SEC3-GGXX01117148.1
Chabr1-SEC3-GGXX01114400.1
Nitmi2-SEC3-JV802182.772660.800864.SRR
Nitmi1-SEC3-JV812537.1
Charophyceae
Colir-SEC3-QPDY_2006926
Color-SEC3-GBSL01050663.1
Coleochaetophyceae
Kleni-SEC3-kfl00782_0020_v1.1
Intpa-SEC3-FPCO_2005353.ERR2041090.2914674.1
Strca-SEC3-Scap_050567
Klebsormidiophyceae
Spimi-SEC3-NNHQ_2006744
Chlat-SEC3-scaffold409.fgen
Mesvi-SEC3-GBSK01006747.01040386.1
Mesostigmatophyceae
Chlamydomonas_reinhardtii-SEC3-Cre09.g392000.fgen
Volvox_carteri-SEC3-XM_002947564.1
Chromochloris_zofingiensis-SEC3-Cz16g15080.t1
Chlorella_sp-SEC3-GIKO01080366.01052845.1
Chlorella_sorokiniana-SEC3-GAPD01039464.01039465.01033088.1
Coccomyxa_subellipsoidea-SEC3-46702.fgen
Chlorophyta
Danio_rerio-EXOC1-A3KPP9
Homo_sapiens-EXOC1-Q9NV70
Drosophila_melanogaster-EXOC1-Q9VVG4
Saccharomyces_cerevisiae-Sec3-P33332
Opisthokonta

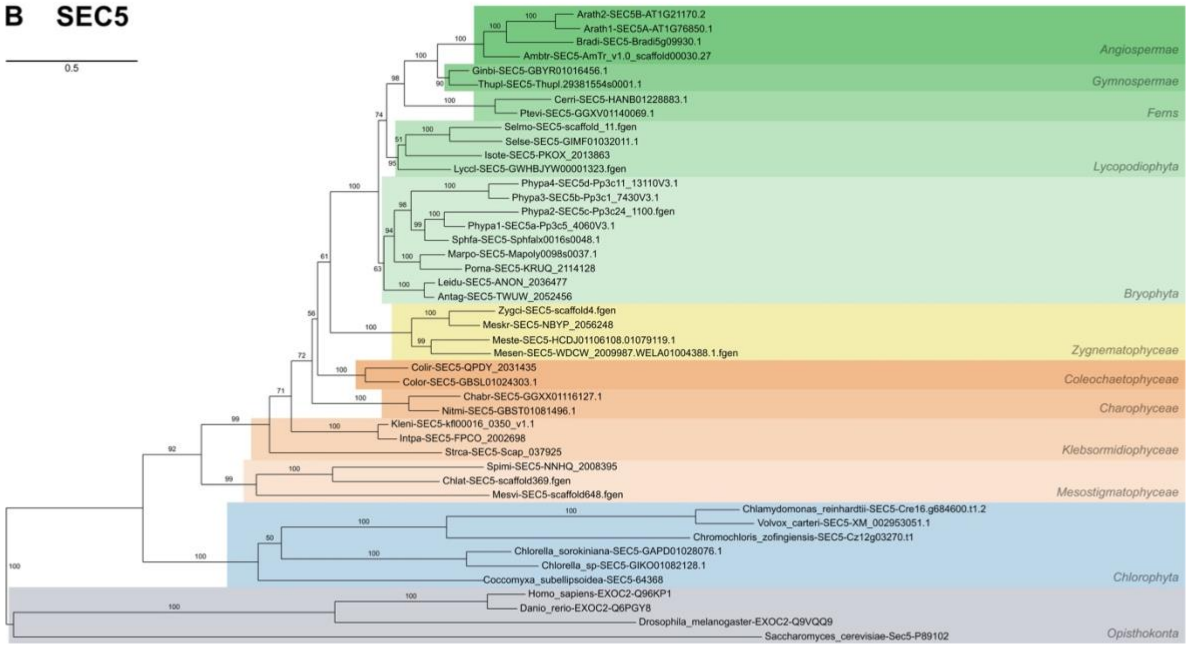
B SEC5
0.5
Arath2-SEC5B-AT1G21170.2
Arath1-SEC5A-AT1G76850.1
Bradi-SEC5-Bradi5g09930.1
Ambtr-SEC5-AmTr_v1.0_scaffold00030.27
Angiospermae
Ginbi-SEC5-GBYR01016456.1
Thupl-SEC5-Thupl.29381554s0001.1
Gymnospermae
Cerri-SEC5-HANB01228883.1
Ptevi-SEC5-GGXV01140069.1
Ferns
Selmo-SEC5-scaffold_11.fgen
Selse-SEC5-GIMF01032011.1
Isote-SEC5-PKOX_2013863
Lyccl-SEC5-GWHBJYW00001323.fgen
Lycopodiophyta
Phypa4-SEC5d-Pp3c11_13110V3.1
Phypa3-SEC5b-Pp3c1_7430V3.1
Phypa2-SEC5c-Pp3c24_1100.fgen
Phypa1-SEC5a-Pp3c5_4060V3.1
Sphfa-SEC5-Sphfalx0016s0048.1
Marpo-SEC5-Mapoly0098s0037.1
Porna-SEC5-KRUQ_2114128
Leidu-SEC5-ANON_2036477
Antag-SEC5-TWUW_2052456
Bryophyta
Zygci-SEC5-scaffold4.fgen
Meskr-SEC5-NBYP_2056248
Meste-SEC5-HCDJ01106108.01079119.1
Mesen-SEC5-WDCW_2009987.WELA01004388.1.fgen
Zygnematophyceae
Colir-SEC5-QPDY_2031435
Color-SEC5-GBSL01024303.1
Coleochaetophyceae
Chabr-SEC5-GGXX01116127.1
Nitmi-SEC5-GBST01081496.1
Charophyceae
Kleni-SEC5-kfl00016_0350_v1.1
Intpa-SEC5-FPCO_2002698
Strca-SEC5-Scap_037925
Klebsormidiophyceae
Spimi-SEC5-NNHQ_2008395
Chlat-SEC5-scaffold369.fgen
Mesvi-SEC5-scaffold648.fgen
Mesostigmatophyceae
Chlamydomonas_reinhardtii-SEC5-Cre16.g684600.t1.2
Volvox_carteri-SEC5-XM_002953051.1
Chromochloris_zofingiensis-SEC5-Cz12g03270.t1
Chlorella_sorokiniana-SEC5-GAPD01028076.1
Chlorella_sp-SEC5-GIKO01082128.1
Coccomyxa_subellipsoidea-SEC5-64368
Chlorophyta
Homo_sapiens-EXOC2-Q96KP1
Danio_rerio-EXOC2-Q6PGY8
Drosophila_melanogaster-EXOC2-Q9VQQ9
Saccharomyces_cerevisiae-Sec5-P89102
Opisthokonta

**C SEC6**

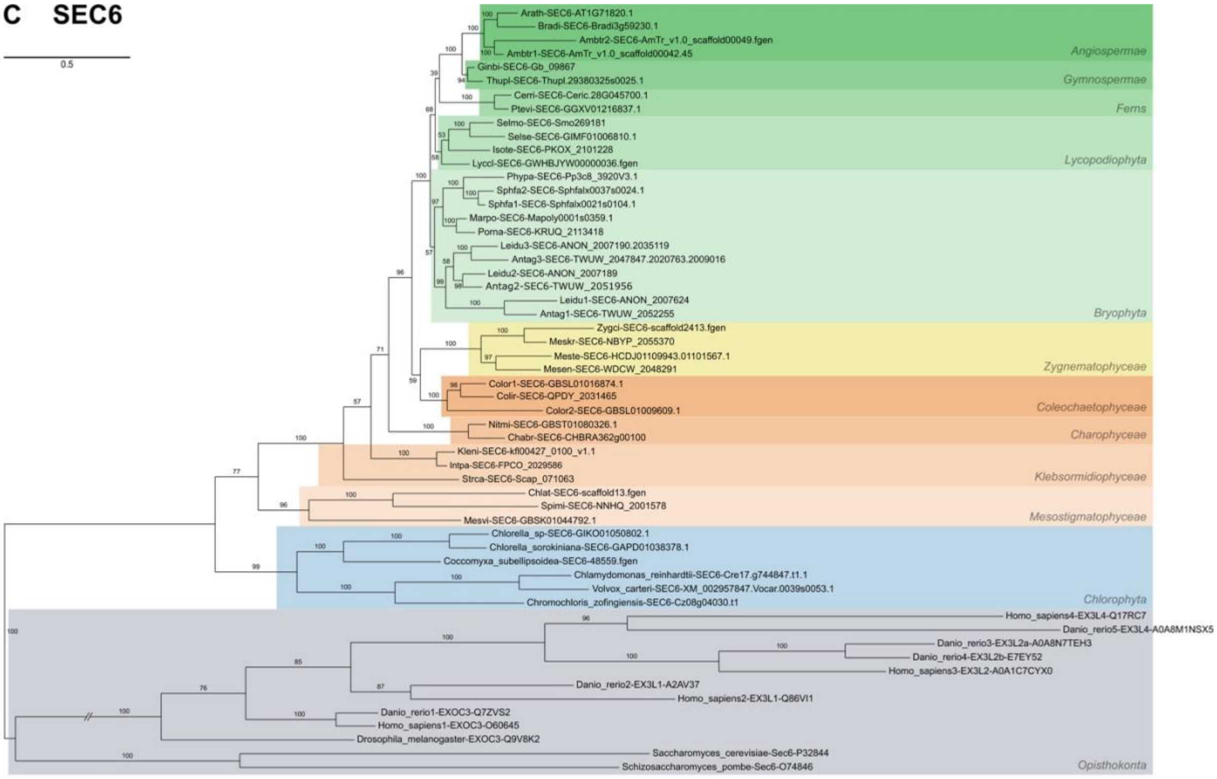
0.5
Arath-SEC6-AT1G71820.1
Bradi-SEC6-Bradi3g59230.1
Ambtr2-SEC6-AmTr_v1.0_scaffold00049.fgen
Ambtr1-SEC6-AmTr_v1.0_scaffold00042.45
Angiospermae
Ginbi-SEC6-Gb_09867
Thupl-SEC6-Thupl.29380325s0025.1
Gymnospermae
Cerri-SEC6-Ceric.28G045700.1
Ptevi-SEC6-GGXV01216837.1
Ferns
Selmo-SEC6-Smo269181
Selse-SEC6-GIMF01006810.1
Isote-SEC6-PKOX_2101228
Lyccl-SEC6-GWHBJYW00000036.fgen
Lycopodiophyta
Phypa-SEC6-Pp3c8_3920V3.1
Sphfa2-SEC6-Sphfalx0037s0024.1
Sphfa1-SEC6-Sphfalx0021s0104.1
Marpo-SEC6-Mapoly0001s0359.1
Porna-SEC6-KRUQ_2113418
Leidu3-SEC6-ANON_2007190.2035119
Antag3-SEC6-TWUW_2047847.2020763.2009016
Leidu2-SEC6-ANON_2007189
Antag2-SEC6-TWUW_2051956
Leidu1-SEC6-ANON_2007624
Antag1-SEC6-TWUW_2052255
Bryophyta
Zygci-SEC6-scaffold2413.fgen
Meskr-SEC6-NBYP_2055370
Meste-SEC6-HCDJ01109943.01101567.1
Mesen-SEC6-WDCW_2048291
Zygnematophyceae
Color1-SEC6-GBSL01016874.1
Colir-SEC6-QPDY_2031465
Color2-SEC6-GBSL01009609.1
Coleochaetophyceae
Nitmi-SEC6-GBST01080326.1
Chabr-SEC6-CHBRA362g00100
Charophyceae
Kleni-SEC6-kfl00427_0100_v1.1
Intpa-SEC6-FPCO_2029586
Strca-SEC6-Scap_071063
Klebsormidiophyceae
Chlat-SEC6-scaffold13.fgen
Spimi-SEC6-NNHQ_2001578
Mesvi-SEC6-GBSK01044792.1
Mesostigmatophyceae
Chlorella_sp-SEC6-GIKO01050802.1
Chlorella_sorokiniana-SEC6-GAPD01038378.1
Coccomyxa_subellipsoidea-SEC6-48559.fgen
Chlamydomonas_reinhardtii-SEC6-Cre17.g744847.t1.1
Volvox_carteri-SEC6-XM_002957847.Vocar.0039s0053.1
Chromochloris_zofingiensis-SEC6-Cz08g04030.t1
Chlorophyta
Homo_sapiens4-EX3L4-Q17RC7
Danio_rerio5-EX3L4-A0A8M1NSX5
Danio_rerio3-EX3L2a-A0A8N7TEH3
Danio_rerio4-EX3L2b-E7EY52
Homo_sapiens3-EX3L2-A0A1C7CYX0
Danio_rerio2-EX3L1-A2AV37
Homo_sapiens2-EX3L1-Q86VI1
Danio_rerio1-EXOC3-Q7ZVS2
Homo_sapiens1-EXOC3-O60645
Drosophila_melanogaster-EXOC3-Q9V8K2
Saccharomyces_cerevisiae-Sec6-P32844
Schizosaccharomyces_pombe-Sec6-O74846
Opisthokonta

**D SEC8**

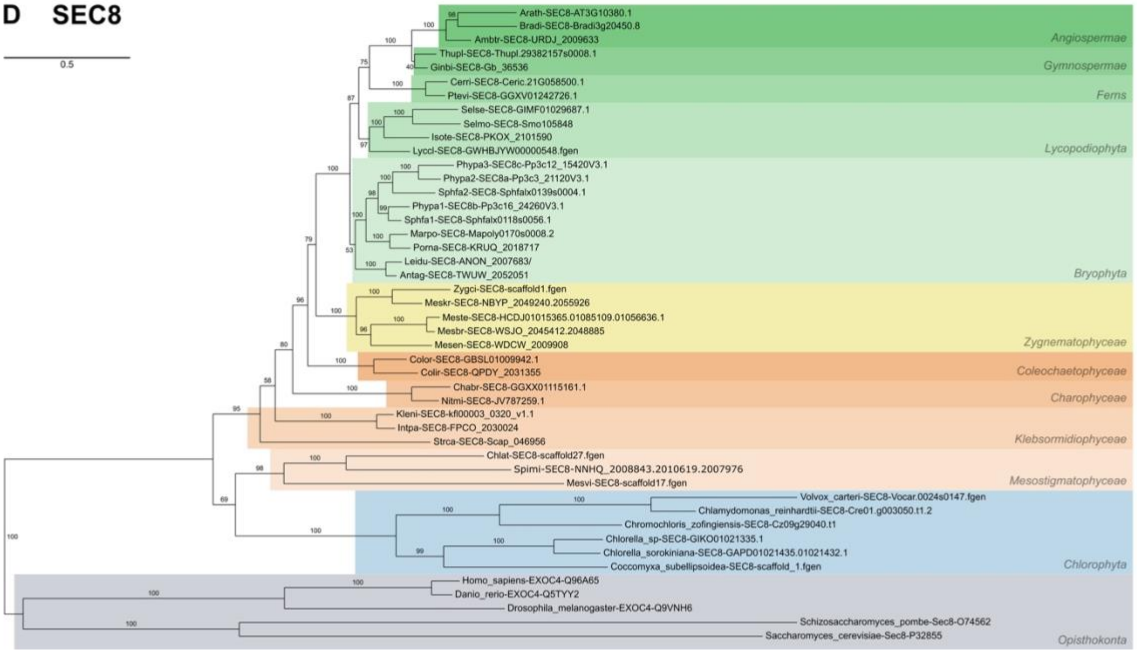
0.5
Arath-SEC8-AT3G10380.1
Bradi-SEC8-Bradi3g20450.8
Ambtr-SEC8-URDJ_2009633
Angiospermae
Thupl-SEC8-Thupl.29382157s0008.1
Ginbi-SEC8-Gb_36536
Gymnospermae
Cerri-SEC8-Ceric.21G058500.1
Ptevi-SEC8-GGXV01242726.1
Ferns
Selse-SEC8-GIMF01029687.1
Selmo-SEC8-Smo105848
Isote-SEC8-PKOX_2101590
Lyccl-SEC8-GWHBJYW00000548.fgen
Lycopodiophyta
Phypa3-SEC8c-Pp3c12_15420V3.1
Phypa2-SEC8a-Pp3c3_21120V3.1
Sphfa2-SEC8-Sphfalx0139s0004.1
Phypa1-SEC8b-Pp3c16_24260V3.1
Sphfa1-SEC8-Sphfalx0118s0056.1
Marpo-SEC8-Mapoly0170s0008.2
Porna-SEC8-KRUQ_2018717
Leidu-SEC8-ANON_2007683/
Antag-SEC8-TWUW_2052051
Bryophyta
Zygci-SEC8-scaffold1.fgen
Meskr-SEC8-NBYP_2049240.2055926
Meste-SEC8-HCDJ01015365.01085109.01056636.1
Mesbr-SEC8-WSJO_2045412.2048885
Mesen-SEC8-WDCW_2009908
Zygnematophyceae
Color-SEC8-GBSL01009942.1
Colir-SEC8-QPDY_2031355
Coleochaetophyceae
Chabr-SEC8-GGXX01115161.1
Nitmi-SEC8-JV787259.1
Charophyceae
Kleni-SEC8-kfl00003_0320_v1.1
Intpa-SEC8-FPCO_2030024
Strca-SEC8-Scap_046956
Klebsormidiophyceae
Chlat-SEC8-scaffold27.fgen
Spimi-SEC8-NNHQ_2008843.2010619.2007976
Mesvi-SEC8-scaffold17.fgen
Mesostigmatophyceae
Volvox_carteri-SEC8-Vocar.0024s0147.fgen
Chlamydomonas_reinhardtii-SEC8-Cre01.g003050.t1.2
Chromochloris_zofingiensis-SEC8-Cz09g29040.t1
Chlorella_sp-SEC8-GIKO01021335.1
Chlorella_sorokiniana-SEC8-GAPD01021435.01021432.1
Coccomyxa_subellipsoidea-SEC8-scaffold_1.fgen
Chlorophyta
Homo_sapiens-EXOC4-Q96A65
Danio_rerio-EXOC4-Q5TYY2
Drosophila_melanogaster-EXOC4-Q9VNH6
Schizosaccharomyces_pombe-Sec8-O74562
Saccharomyces_cerevisiae-Sec8-P32855
Opisthokonta

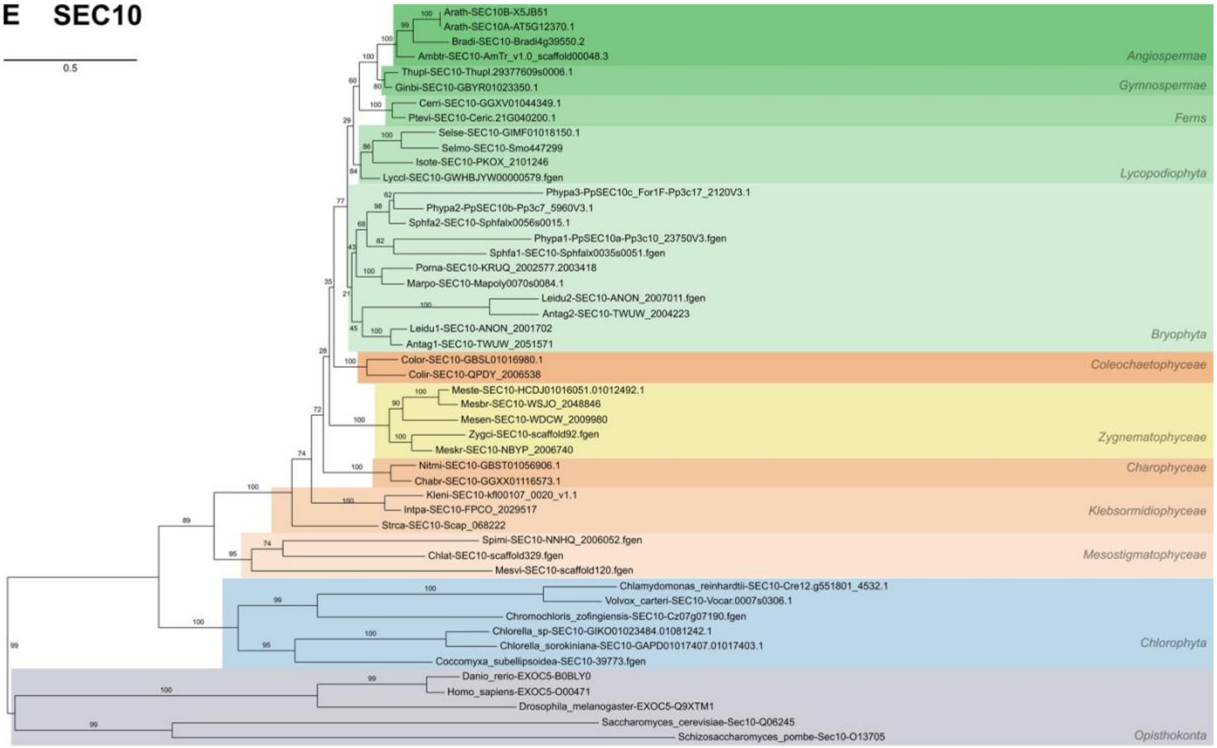
E SEC10
0.5
Arath-SEC10B-X5JB51
Arath-SEC10A-AT5G12370.1
Bradi-SEC10-Bradi4g39550.2
Ambtr-SEC10-AmTr_v1.0_scaffold00048.3
Angiospermae
Thupl-SEC10-Thupl.29377609s0006.1
Ginbi-SEC10-GBYR01023350.1
Gymnospermae
Cerri-SEC10-GGXV01044349.1
Ptevi-SEC10-Ceric.21G040200.1
Ferns
Selse-SEC10-GIMF01018150.1
Selmo-SEC10-Smo447299
Isote-SEC10-PKOX_2101246
Lyccl-SEC10-GWHBJYW00000579.fgen
Lycopodiophyta
Phypa3-PpSEC10c_For1F-Pp3c17_2120V3.1
Phypa2-PpSEC10b-Pp3c7_5960V3.1
Sphfa2-SEC10-Sphfalx0056s0015.1
Phypa1-PpSEC10a-Pp3c10_23750V3.fgen
Sphfa1-SEC10-Sphfalx0035s0051.fgen
Porna-SEC10-KRUQ_2002577.2003418
Marpo-SEC10-Mapoly0070s0084.1
Leidu2-SEC10-ANON_2007011.fgen
Antag2-SEC10-TWUW_2004223
Leidu1-SEC10-ANON_2001702
Antag1-SEC10-TWUW_2051571
Bryophyta
Color-SEC10-GBSL01016980.1
Colir-SEC10-QPDY_2006538
Coleochaetophyceae
Meste-SEC10-HCDJ01016051.01012492.1
Mesbr-SEC10-WSJO_2048846
Mesen-SEC10-WDCW_2009980
Zygci-SEC10-scaffold92.fgen
Meskr-SEC10-NBYP_2006740
Zygnematophyceae
Nitmi-SEC10-GBST01056906.1
Chabr-SEC10-GGXX01116573.1
Charophyceae
Kleni-SEC10-kfl00107_0020_v1.1
Intpa-SEC10-FPCO_2029517
Strca-SEC10-Scap_068222
Klebsormidiophyceae
Spimi-SEC10-NNHQ_2006052.fgen
Chlat-SEC10-scaffold329.fgen
Mesvi-SEC10-scaffold120.fgen
Mesostigmatophyceae
Chlamydomonas_reinhardtii-SEC10-Cre12.g551801_4532.1
Volvox_carteri-SEC10-Vocar.0007s0306.1
Chromochloris_zofingiensis-SEC10-Cz07g07190.fgen
Chlorella_sp-SEC10-GIKO01023484.01081242.1
Chlorella_sorokiniana-SEC10-GAPD01017407.01017403.1
Coccomyxa_subellipsoidea-SEC10-39773.fgen
Chlorophyta
Danio_rerio-EXOC5-B0BLY0
Homo_sapiens-EXOC5-O00471
Drosophila_melanogaster-EXOC5-Q9XTM1
Saccharomyces_cerevisiae-Sec10-Q06245
Schizosaccharomyces_pombe-Sec10-O13705
Opisthokonta

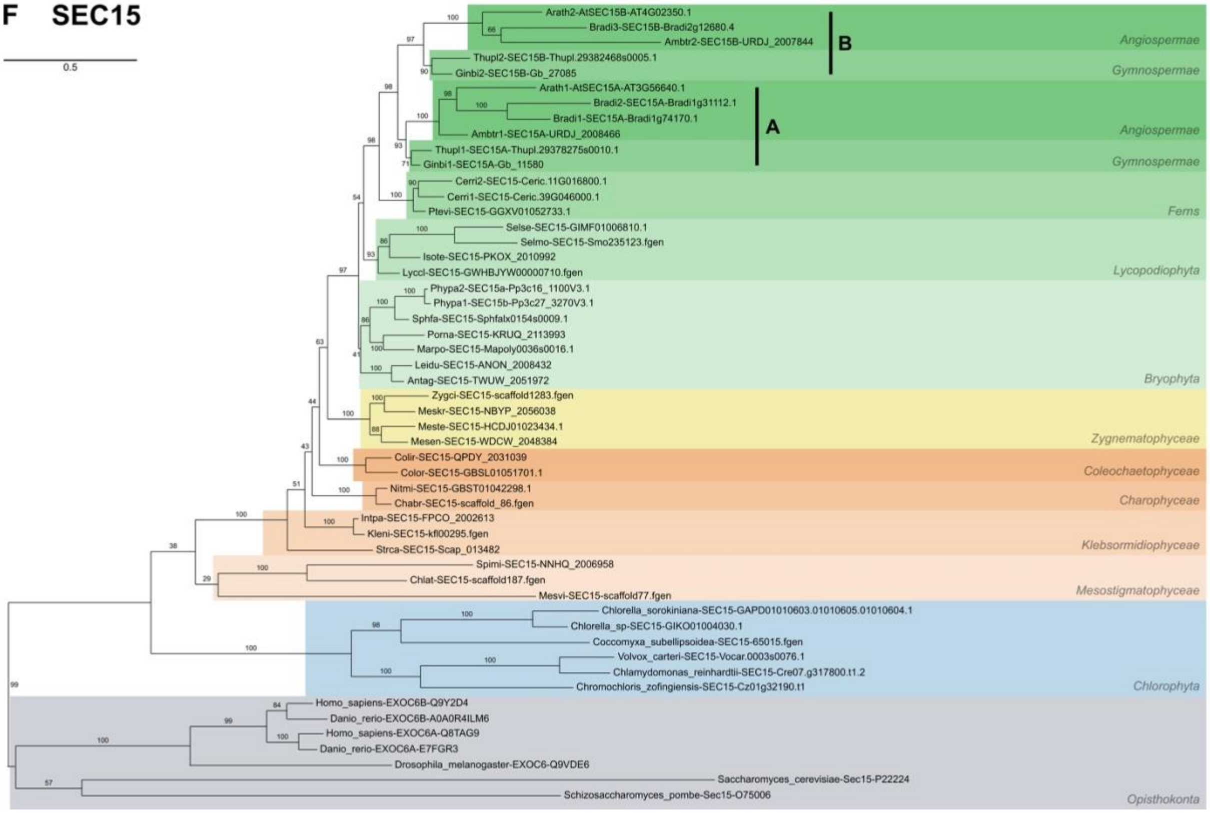
F SEC15
0.5
Arath2-AtSEC15B-AT4G02350.1
Bradi3-SEC15B-Bradi2g12680.4
Ambtr2-SEC15B-URDJ_2007844
B
Angiospermae
Thupl2-SEC15B-Thupl.29382468s0005.1
Ginbi2-SEC15B-Gb_27085
Gymnospermae
Arath1-AtSEC15A-AT3G56640.1
Bradi2-SEC15A-Bradi1g31112.1
Bradi1-SEC15A-Bradi1g74170.1
Ambtr1-SEC15A-URDJ_2008466
A
Angiospermae
Thupl1-SEC15A-Thupl.29378275s0010.1
Ginbi1-SEC15A-Gb_11580
Gymnospermae
Cerri2-SEC15-Ceric.11G016800.1
Cerri1-SEC15-Ceric.39G046000.1
Ptevi-SEC15-GGXV01052733.1
Ferns
Selse-SEC15-GIMF01006810.1
Selmo-SEC15-Smo235123.fgen
Isote-SEC15-PKOX_2010992
Lyccl-SEC15-GWHBJYW00000710.fgen
Lycopodiophyta
Phypa2-SEC15a-Pp3c16_1100V3.1
Phypa1-SEC15b-Pp3c27_3270V3.1
Sphfa-SEC15-Sphfalx0154s0009.1
Porna-SEC15-KRUQ_2113993
Marpo-SEC15-Mapoly0036s0016.1
Leidu-SEC15-ANON_2008432
Antag-SEC15-TWUW_2051972
Bryophyta
Zygci-SEC15-scaffold1283.fgen
Meskr-SEC15-NBYP_2056038
Meste-SEC15-HCDJ01023434.1
Mesen-SEC15-WDCW_2048384
Zygnematophyceae
Colir-SEC15-QPDY_2031039
Color-SEC15-GBSL01051701.1
Coleochaetophyceae
Nitmi-SEC15-GBST01042298.1
Chabr-SEC15-scaffold_86.fgen
Charophyceae
Intpa-SEC15-FPCO_2002613
Kleni-SEC15-kfl00295.fgen
Strca-SEC15-Scap_013482
Klebsormidiophyceae
Spimi-SEC15-NNHQ_2006958
Chlat-SEC15-scaffold187.fgen
Mesvi-SEC15-scaffold77.fgen
Mesostigmatophyceae
Chlorella_sorokiniana-SEC15-GAPD01010603.01010605.01010604.1
Chlorella_sp-SEC15-GIKO01004030.1
Coccomyxa_subellipsoidea-SEC15-65015.fgen
Volvox_carteri-SEC15-Vocar.0003s0076.1
Chlamydomonas_reinhardtii-SEC15-Cre07.g317800.t1.2
Chromochloris_zofingiensis-SEC15-Cz01g32190.t1
Chlorophyta
Homo_sapiens-EXOC6B-Q9Y2D4
Danio_rerio-EXOC6B-A0A0R4ILM6
Homo_sapiens-EXOC6A-Q8TAG9
Danio_rerio-EXOC6A-E7FGR3
Drosophila_melanogaster-EXOC6-Q9VDE6
Saccharomyces_cerevisiae-Sec15-P22224
Schizosaccharomyces_pombe-Sec15-O75006
Opisthokonta

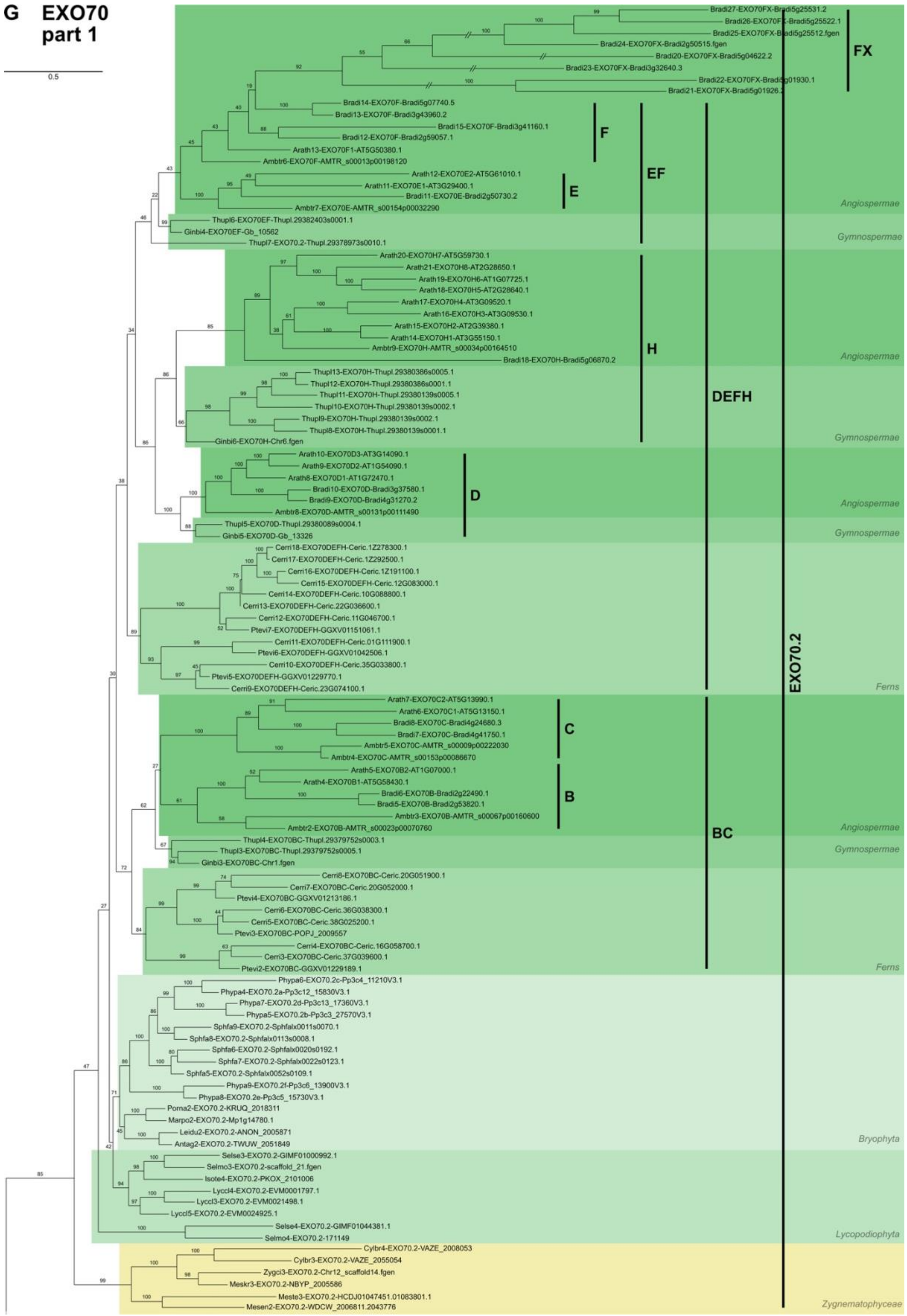

G EXO70
part 1
0.5
Bradi27-EXO70FX-Bradi5g25531.2
Bradi26-EXO70FX-Bradi5g25522.1
Bradi25-EXO70FX-Bradi5g25512.fgen
Bradi24-EXO70FX-Bradi2g50515.fgen
Bradi20-EXO70FX-Bradi5g04622.2
Bradi23-EXO70FX-Bradi3g32640.3
Bradi22-EXO70FX-Bradi5g01930.1
Bradi21-EXO70FX-Bradi5g01926.2
FX
Bradi14-EXO70F-Bradi5g07740.5
Bradi13-EXO70F-Bradi3g43960.2
Bradi15-EXO70F-Bradi3g41160.1
Bradi12-EXO70F-Bradi2g59057.1
Arath13-EXO70F1-AT5G50380.1
Ambtr6-EXO70F-AMTR_s00013p00198120
F
Arath12-EXO70E2-AT5G61010.1
Arath11-EXO70E1-AT3G29400.1
Bradi11-EXO70E-Bradi2g50730.2
Ambtr7-EXO70E-AMTR_s00154p00032290
E
EF
Angiospermae
Thupl6-EXO70EF-Thupl.29382403s0001.1
Ginbi4-EXO70EF-Gb_10562
Thupl7-EXO70.2-Thupl.29378973s0010.1
Gymnospermae
Arath20-EXO70H7-AT5G59730.1
Arath21-EXO70H8-AT2G28650.1
Arath19-EXO70H6-AT1G07725.1
Arath18-EXO70H5-AT2G28640.1
Arath17-EXO70H4-AT3G09520.1
Arath16-EXO70H3-AT3G09530.1
Arath15-EXO70H2-AT2G39380.1
Arath14-EXO70H1-AT3G55150.1
Ambtr9-EXO70H-AMTR_s00034p00164510
Bradi18-EXO70H-Bradi5g06870.2
H
Angiospermae
Thupl13-EXO70H-Thupl.29380386s0005.1
Thupl12-EXO70H-Thupl.29380386s0001.1
Thupl11-EXO70H-Thupl.29380139s0005.1
Thupl10-EXO70H-Thupl.29380139s0002.1
Thupl9-EXO70H-Thupl.29380139s0002.1
Thupl8-EXO70H-Thupl.29380139s0001.1
Ginbi6-EXO70H-Chr6.fgen
DEFH
Gymnospermae
Arath10-EXO70D3-AT3G14090.1
Arath9-EXO70D2-AT1G54090.1
Arath8-EXO70D1-AT1G72470.1
Bradi10-EXO70D-Bradi3g37580.1
Bradi9-EXO70D-Bradi4g31270.2
Ambtr8-EXO70D-AMTR_s00131p00111490
D
Angiospermae
Thupl5-EXO70D-Thupl.29380089s0004.1
Ginbi5-EXO70D-Gb_13326
Gymnospermae
Cerri18-EXO70DEFH-Ceric.1Z278300.1
Cerri17-EXO70DEFH-Ceric.1Z292500.1
Cerri16-EXO70DEFH-Ceric.1Z191100.1
Cerri15-EXO70DEFH-Ceric.12G083000.1
Cerri14-EXO70DEFH-Ceric.10G088800.1
Cerri13-EXO70DEFH-Ceric.22G036600.1
Cerri12-EXO70DEFH-Ceric.11G046700.1
Ptevi7-EXO70DEFH-GGXV01151061.1
Cerri11-EXO70DEFH-Ceric.01G111900.1
Ptevi6-EXO70DEFH-GGXV01042506.1
Cerri10-EXO70DEFH-Ceric.35G033800.1
Ptevi5-EXO70DEFH-GGXV01229770.1
Cerri9-EXO70DEFH-Ceric.23G074100.1
EXO70.2
Ferns
Arath7-EXO70C2-AT5G13990.1
Arath6-EXO70C1-AT5G13150.1
Bradi8-EXO70C-Bradi4g24680.3
Bradi7-EXO70C-Bradi4g41750.1
Ambtr5-EXO70C-AMTR_s00009p00222030
Ambtr4-EXO70C-AMTR_s00153p00086670
C
Arath5-EXO70B2-AT1G07000.1
Arath4-EXO70B1-AT5G58430.1
Bradi6-EXO70B-Bradi2g22490.1
Bradi5-EXO70B-Bradi2g53820.1
Ambtr3-EXO70B-AMTR_s00067p00160600
Ambtr2-EXO70B-AMTR_s00023p00070760
B
BC
Angiospermae
Thupl4-EXO70BC-Thupl.29379752s0003.1
Thupl3-EXO70BC-Thupl.29379752s0005.1
Ginbi3-EXO70BC-Chr1.fgen
Gymnospermae
Cerri8-EXO70BC-Ceric.20G051900.1
Cerri7-EXO70BC-Ceric.20G052000.1
Ptevi4-EXO70BC-GGXV01213186.1
Cerri6-EXO70BC-Ceric.36G038300.1
Cerri5-EXO70BC-Ceric.38G025200.1
Ptevi3-EXO70BC-POPJ_2009557
Cerri4-EXO70BC-Ceric.16G058700.1
Cerri3-EXO70BC-Ceric.37G039600.1
Ptevi2-EXO70BC-GGXV01229189.1
Ferns
Phypa6-EXO70.2c-Pp3c4_11210V3.1
Phypa4-EXO70.2a-Pp3c12_15830V3.1
Phypa7-EXO70.2d-Pp3c13_17360V3.1
Phypa5-EXO70.2b-Pp3c3_27570V3.1
Sphfa9-EXO70.2-Sphfalx0011s0070.1
Sphfa8-EXO70.2-Sphfalx0113s0008.1
Sphfa6-EXO70.2-Sphfalx0020s0192.1
Sphfa7-EXO70.2-Sphfalx0022s0123.1
Sphfa5-EXO70.2-Sphfalx0052s0109.1
Phypa9-EXO70.2f-Pp3c6_13900V3.1
Phypa8-EXO70.2e-Pp3c5_15730V3.1
Porna2-EXO70.2-KRUQ_2018311
Marpo2-EXO70.2-Mp1g14780.1
Leidu2-EXO70.2-ANON_2005871
Antag2-EXO70.2-TWUW_2051849
Bryophyta
Selse3-EXO70.2-GIMF01000992.1
Selmo3-EXO70.2-scaffold_21.fgen
Isote4-EXO70.2-PKOX_2101006
Lyccl4-EXO70.2-EVM0001797.1
Lyccl3-EXO70.2-EVM0021498.1
Lyccl5-EXO70.2-EVM0024925.1
Selse4-EXO70.2-GIMF01044381.1
Selmo4-EXO70.2-171149
Lycopodiophyta
Cylbr4-EXO70.2-VAZE_2008053
Cylbr3-EXO70.2-VAZE_2055054
Zygci3-EXO70.2-Chr12_scaffold14.fgen
Meskr3-EXO70.2-NBYP_2005586
Meste3-EXO70.2-HCDJ01047451.01083801.1
Mesen2-EXO70.2-WDCW_2006811.2043776
Zygnematophyceae

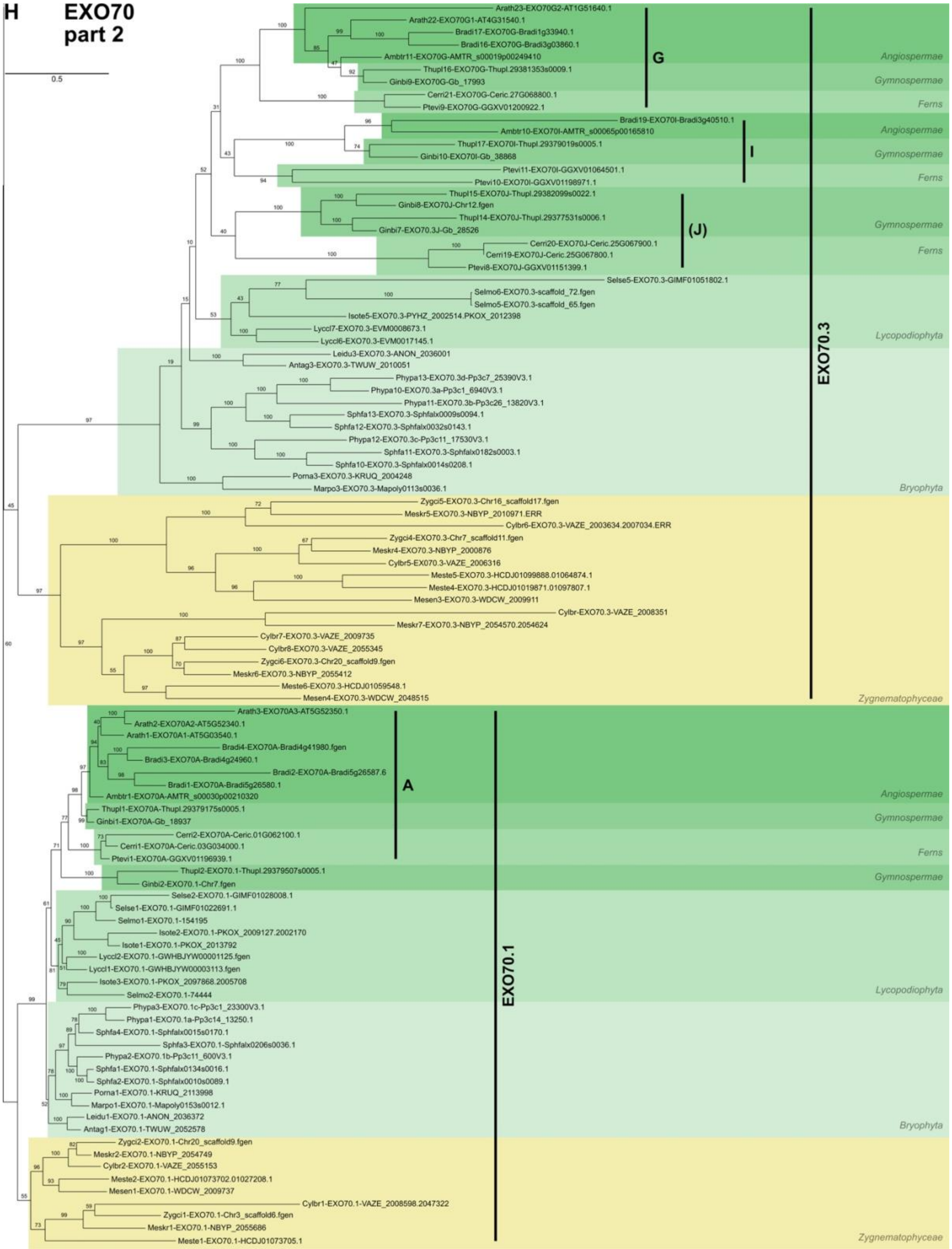
H
EXO70
part 2
0.5
Arath23-EXO70G2-AT1G51640.1
Arath22-EXO70G1-AT4G31540.1
Bradi17-EXO70G-Bradi1g33940.1
Bradi16-EXO70G-Bradi3g03860.1
Ambtr11-EXO70G-AMTR_s00019p00249410
Thupl16-EXO70G-Thupl.29381353s0009.1
Ginbi9-EXO70G-Gb_17993
Cerri21-EXO70G-Ceric.27G068800.1
Ptevi9-EXO70G-GGXV01200922.1
G
Bradi19-EXO70I-Bradi3g40510.1
Ambtr10-EXO70I-AMTR_s00065p00165810
Thupl17-EXO70I-Thupl.29379019s0005.1
Ginbi10-EXO70I-Gb_38868
Ptevi11-EXO70I-GGXV01064501.1
Ptevi10-EXO70I-GGXV01198971.1
I
Thupl15-EXO70J-Thupl.29382099s0022.1
Ginbi8-EXO70J-Chr12.fgen
Thupl14-EXO70J-Thupl.29377531s0006.1
Ginbi7-EXO70.3J-Gb_28526
Cerri20-EXO70J-Ceric.25G067900.1
Cerri19-EXO70J-Ceric.25G067800.1
Ptevi8-EXO70J-GGXV01151399.1
(J)
Selse5-EXO70.3-GIMF01051802.1
Selmo6-EXO70.3-scaffold_72.fgen
Selmo5-EXO70.3-scaffold_65.fgen
Isote5-EXO70.3-PYHZ_2002514.PKOX_2012398
Lyccl7-EXO70.3-EVM0008673.1
Lyccl6-EXO70.3-EVM0017145.1
Leidu3-EXO70.3-ANON_2036001
Antag3-EXO70.3-TWUW_2010051
Phypa13-EXO70.3d-Pp3c7_25390V3.1
Phypa10-EXO70.3a-Pp3c1_6940V3.1
Phypa11-EXO70.3b-Pp3c26_13820V3.1
Sphfa13-EXO70.3-Sphfalx0009s0094.1
Sphfa12-EXO70.3-Sphfalx0032s0143.1
Phypa12-EXO70.3c-Pp3c11_17530V3.1
Sphfa11-EXO70.3-Sphfalx0182s0003.1
Sphfa10-EXO70.3-Sphfalx0014s0208.1
Porna3-EXO70.3-KRUQ_2004248
Marpo3-EXO70.3-Mapoly0113s0036.1
Zygci5-EXO70.3-Chr16_scaffold17.fgen
Meskr5-EXO70.3-NBYP_2010971.ERR
Cylbr6-EXO70.3-VAZE_2003634.2007034.ERR
Zygci4-EXO70.3-Chr7_scaffold11.fgen
Meskr4-EXO70.3-NBYP_2000876
Cylbr5-EXO70.3-VAZE_2006316
Meste5-EXO70.3-HCDJ01099888.01064874.1
Meste4-EXO70.3-HCDJ01019871.01097807.1
Mesen3-EXO70.3-WDCW_2009911
Cylbr-EXO70.3-VAZE_2008351
Meskr7-EXO70.3-NBYP_2054570.2054624
Cylbr7-EXO70.3-VAZE_2009735
Cylbr8-EXO70.3-VAZE_2055345
Zygci6-EXO70.3-Chr20_scaffold9.fgen
Meskr6-EXO70.3-NBYP_2055412
Meste6-EXO70.3-HCDJ01059548.1
Mesen4-EXO70.3-WDCW_2048515
EXO70.3
Arath3-EXO70A3-AT5G52350.1
Arath2-EXO70A2-AT5G52340.1
Arath1-EXO70A1-AT5G03540.1
Bradi4-EXO70A-Bradi4g41980.fgen
Bradi3-EXO70A-Bradi4g24960.1
Bradi2-EXO70A-Bradi5g26587.6
Bradi1-EXO70A-Bradi5g26580.1
Ambtr1-EXO70A-AMTR_s00030p00210320
Thupl1-EXO70A-Thupl.29379175s0005.1
Ginbi1-EXO70A-Gb_18937
Cerri2-EXO70A-Ceric.01G062100.1
Cerri1-EXO70A-Ceric.03G034000.1
Ptevi1-EXO70A-GGXV01196939.1
A
Thupl2-EXO70.1-Thupl.29379507s0005.1
Ginbi2-EXO70.1-Chr7.fgen
Selse2-EXO70.1-GIMF01028008.1
Selse1-EXO70.1-GIMF01022691.1
Selmo1-EXO70.1-154195
Isote2-EXO70.1-PKOX_2009127.2002170
Isote1-EXO70.1-PKOX_2013792
Lyccl2-EXO70.1-GWHBJYW00001125.fgen
Lyccl1-EXO70.1-GWHBJYW00003113.fgen
Isote3-EXO70.1-PKOX_2097868.2005708
Selmo2-EXO70.1-74444
Phypa3-EXO70.1c-Pp3c1_23300V3.1
Phypa1-EXO70.1a-Pp3c14_13250.1
Sphfa4-EXO70.1-Sphfalx0015s0170.1
Sphfa3-EXO70.1-Sphfalx0206s0036.1
Phypa2-EXO70.1b-Pp3c11_600V3.1
Sphfa1-EXO70.1-Sphfalx0134s0016.1
Sphfa2-EXO70.1-Sphfalx0010s0089.1
Porna1-EXO70.1-KRUQ_2113998
Marpo1-EXO70.1-Mapoly0153s0012.1
Leidu1-EXO70.1-ANON_2036372
Antag1-EXO70.1-TWUW_2052578
Zygci2-EXO70.1-Chr20_scaffold9.fgen
Meskr2-EXO70.1-NBYP_2054749
Cylbr2-EXO70.1-VAZE_2055153
Meste2-EXO70.1-HCDJ01073702.01027208.1
Mesen1-EXO70.1-WDCW_2009737
Cylbr1-EXO70.1-VAZE_2008598.2047322
Zygci1-EXO70.1-Chr3_scaffold6.fgen
Meskr1-EXO70.1-NBYP_2055686
Meste1-EXO70.1-HCDJ01073705.1
EXO70.1
Angiospermae
Gymnospermae
Ferns
Lycopodiophyta
Bryophyta
Zygnematophyceae

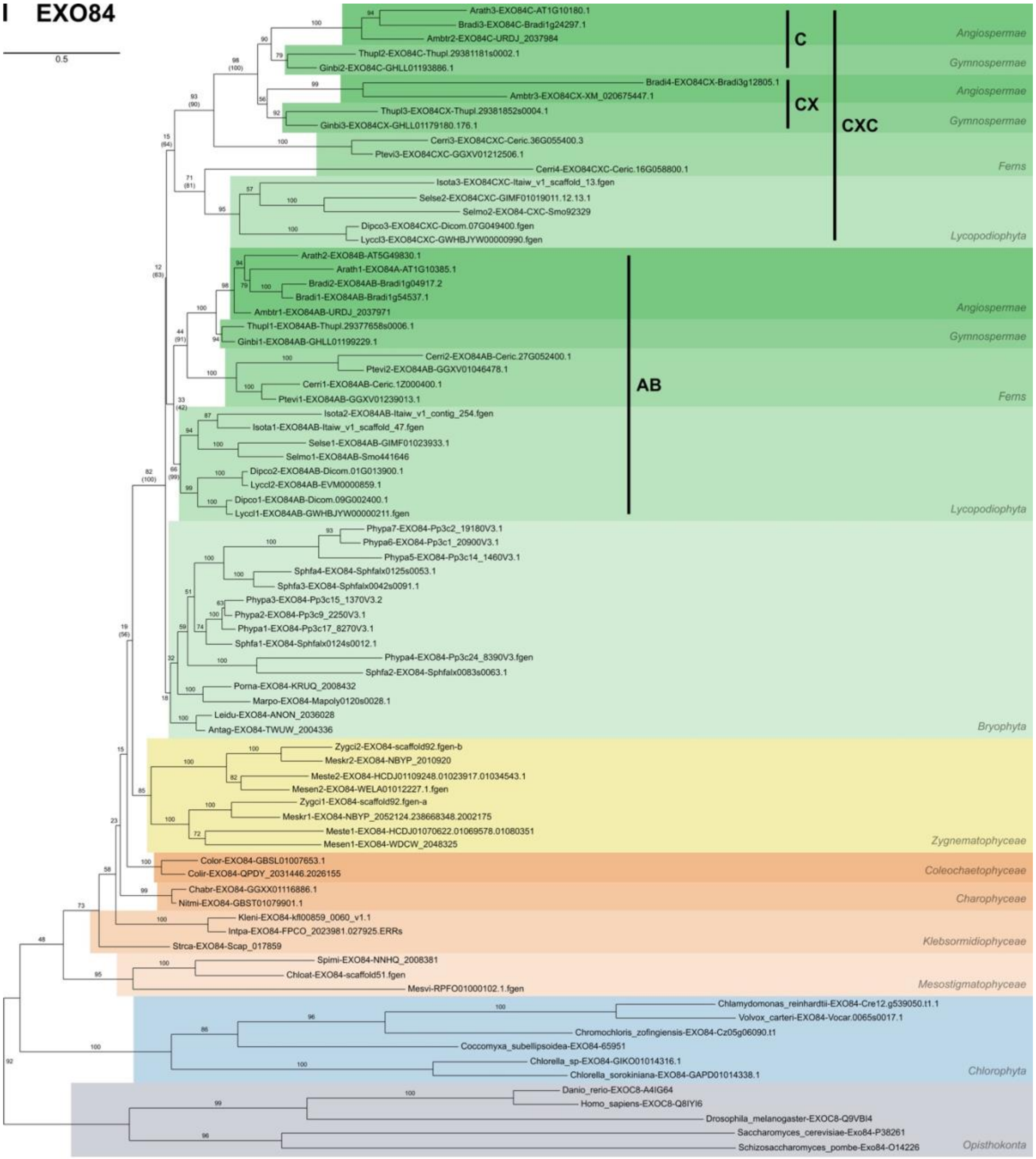


Ambtr – *Amborella trichopoda*, Antag – *Anthoceros agrestis*, Arath – *Arabidopsis thaliana*, Bradi – *Brachypodium distachyon*, Cerri – *Ceratopteris richardii*, Chabr – *Chara braunii*, Chlat – *Chlorokybus atmophyticus*, Colir – *Coleochaete irregularis*, Color – *Coleochaete orbicularis*, Dipco – *Diphasiastrum complanatum*, Ginbi – *Ginkgo biloba*, Intpa – *Interfilum paradoxum*, Isota – *Isoetes taiwanensis*, Isote – *Isoetes tegetiformans*, Kleni – *Klebsormidium nitens*, Leidu – *Leiosporoceros dussii*, Lyccl – *Lycopodium clavatum*, Marpo – *Marchantia polymorpha*, Mesen – *Mesotaenium endlicherianum*, Meskr – *Mesotaenium kramstae*, Meste – *Mesotaenium testaceovaginatum*, Mesvi – *Mesostigma viride*, Nitmi – *Nitella mirabilis*, Phypa – *Physcomitrium patens*, Porna – *Porella navicularis*, Ptevi – *Pteris vittata*, Selmo – *Selaginella moellendorfii*, Selse – *Selaginella sellowii*, Sphfa – *Sphagnum fallax*, Spimi – *Spirotaenia minuta*, Strca – *Streptofilum capillatum*, Thupl – *Thuja plicata*, Zygci – *Zygnema circumcarinatum*.

**Fig. S1. Phylogenetic analyses of SEC3, SEC5, SEC6, SEC8, SEC10, SEC15, EXO70, and EXO84 in streptophytes.** Fulll-length protein sequences were aligned using T-Coffee (SEC3, SEC10), MAFFT (L-INS-i) (SEC5, SEC6, EXO84), or MUSCLE (v3) (SEC8, SEC15, EXO70) methods. Matrices were manually trimmed in combination with Gblocks (SEC3, SEC8, SEC10, SEC15) and/or trimAI (SEC3, EXO84) algorithm outputs. Trees were calculated by NNI (SEC15, EXO84) or NNI+SPR (SEC3, SEC5, SEC6, SEC8, SEC10, EXO70) operations with 500 bootstrap replicates.

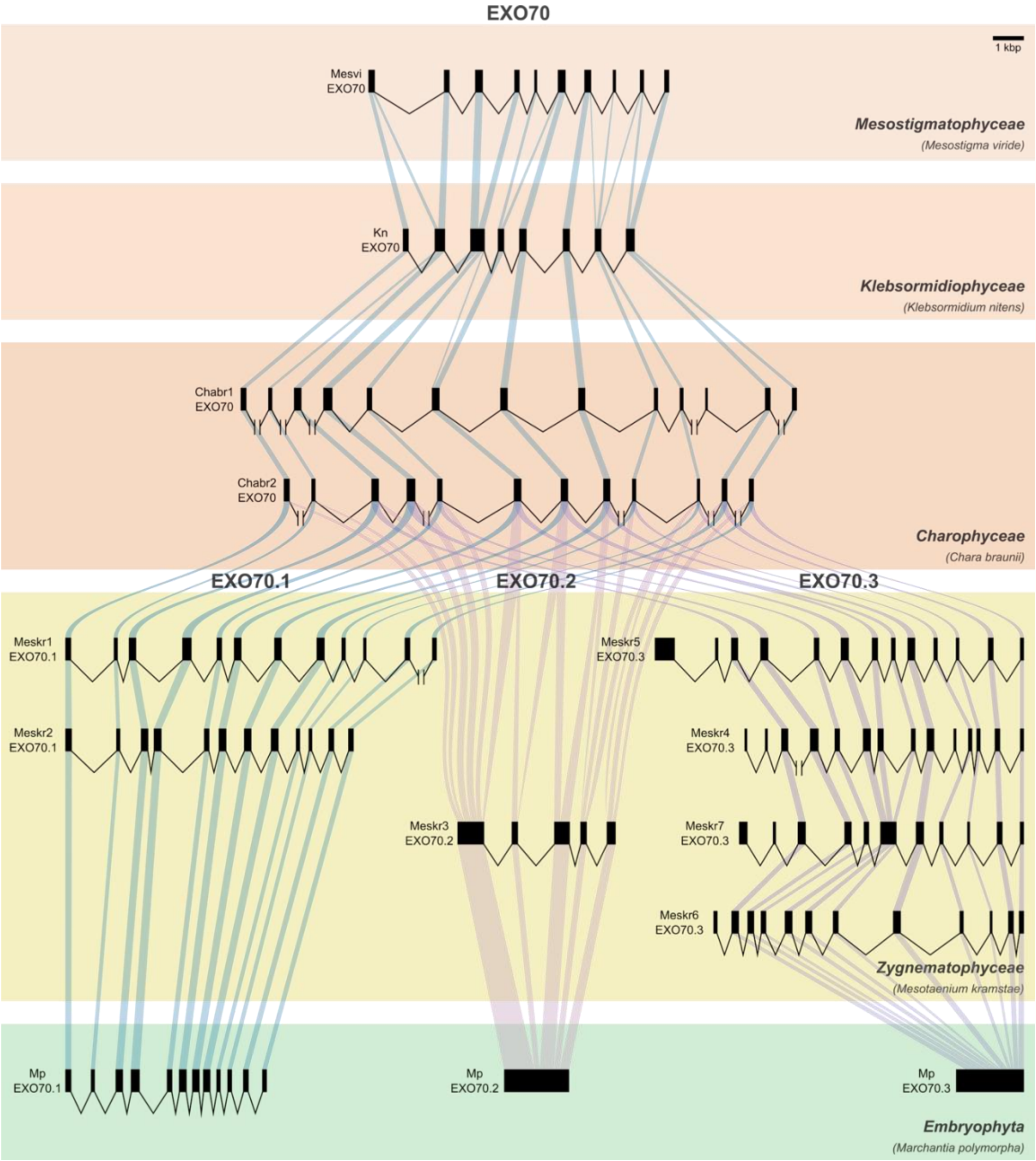


**Fig. S2. Exon–intron architecture of EXO70 homologs from representative streptophyte lineages.** Please note, that only Meskr3-EXO70.2 exons do not share any borders with other EXO70 homologs whatsoever.

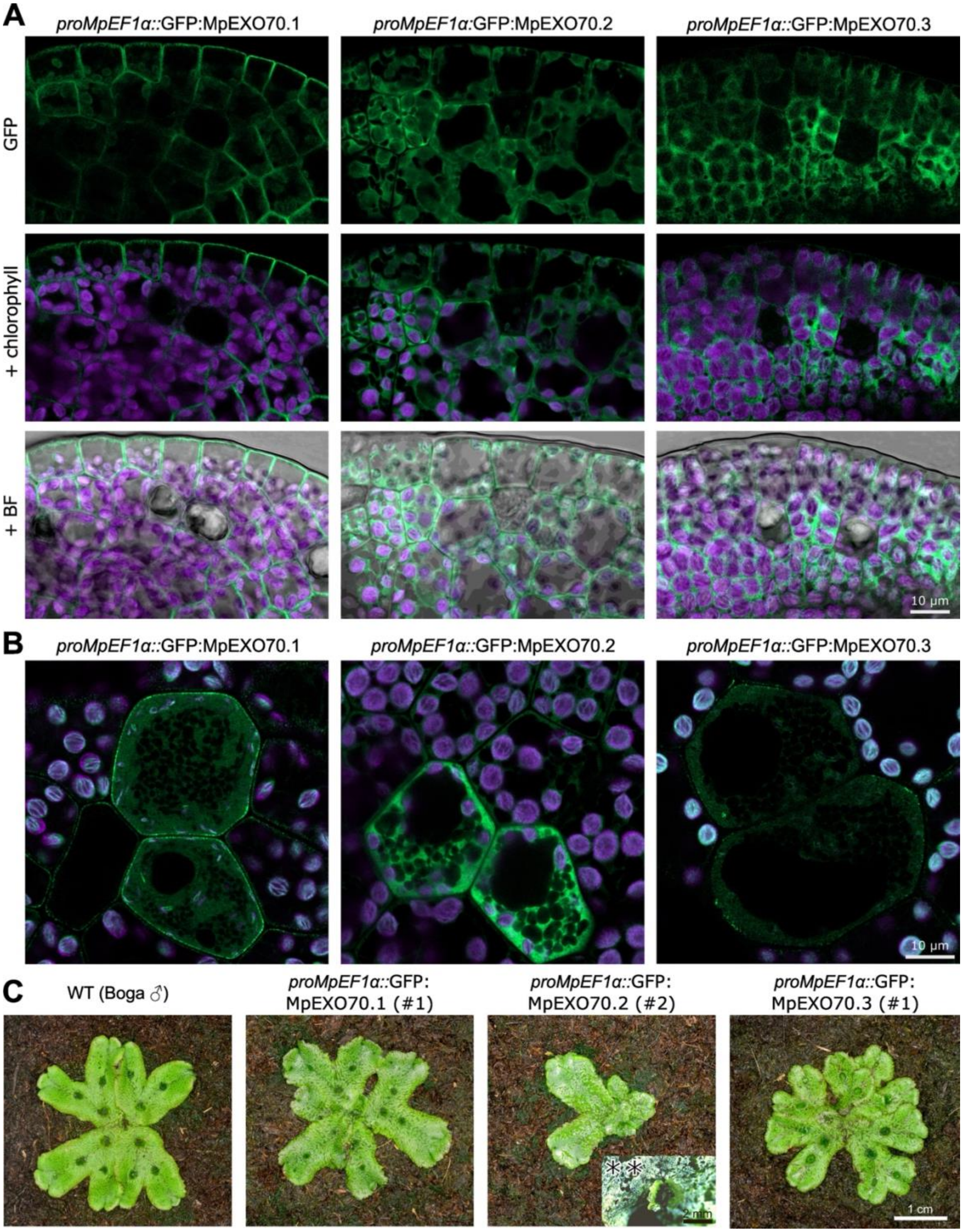


**Fig. S3. Localization and overexpression analyses of Marchantia GFP-tagged EXO70 paralogs in Marchantia gemmae.** Single plane image localization patterns of the tree diverse Marchantia EXO70s in gemmae near the apical meristem (**A**) and initials of the rhizoids (**B**), where *proMpEF1α* is dominantly active. **C)** 5 weeks old thalli grown *ex vitro* (** in *ex vitro* conditions, 8 weeks old thalli over-expressing MpEXO70.2 were able to produce gemmae in malformed gemmae cups).

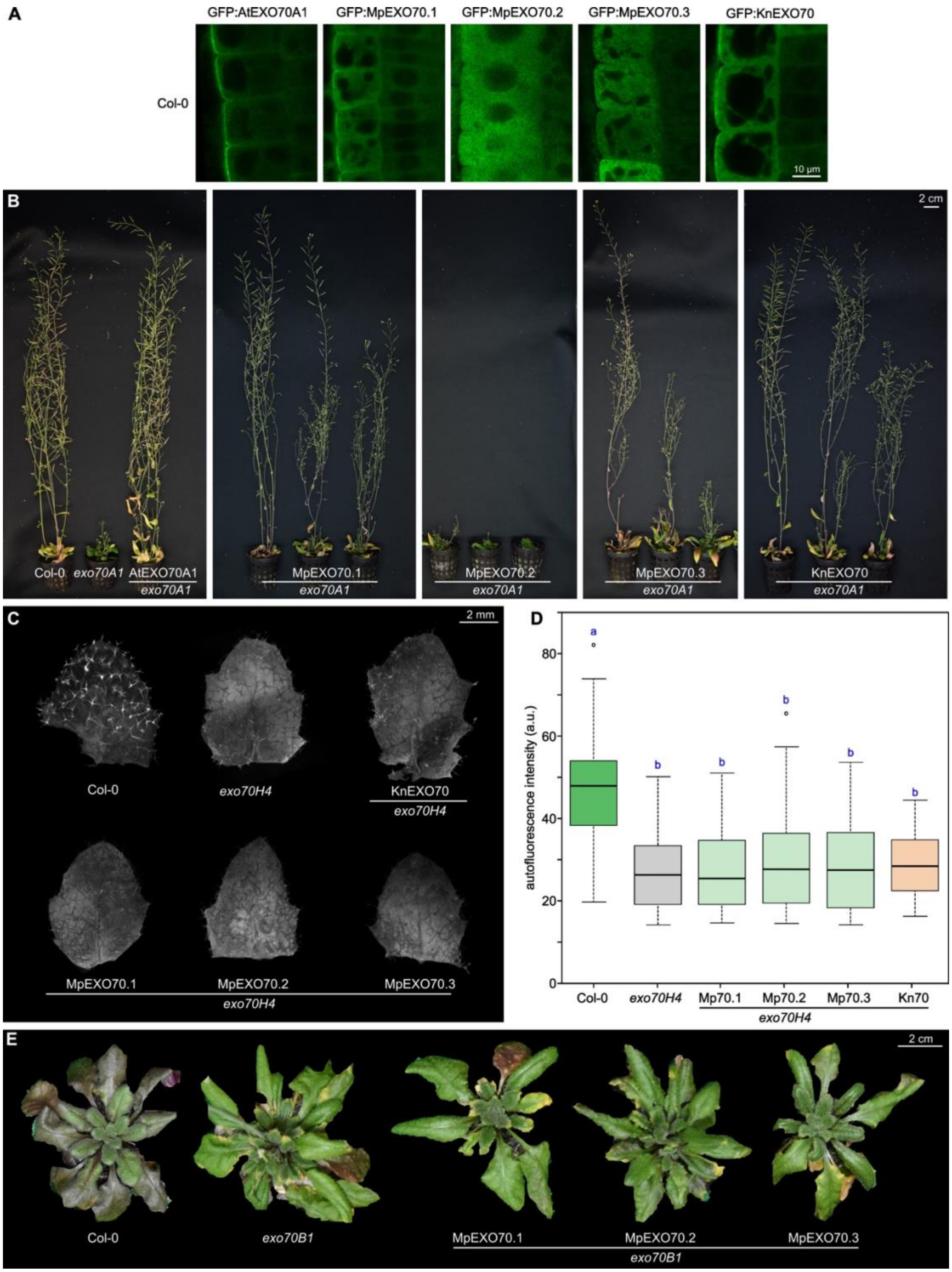


**Fig. S4. Localization, overexpression and complementation analyses of Klebsormidium and Marchantia GFP-tagged EXO70 orthologs in Arabidopsis wild-type, *exo70A1*, *exo70B1* and *exo70H4* mutant backgrounds.** **A** - localization of Arabidopsis EXO70A1, Marchantia EXO70.1-3, and Klebsormidium EXO70 in root epidermal cells of wild type *A. thaliana* (Col-0). **B** - phenotypic variations of 6-week-old plants expressing Marchantia and Klebsormidium EXO70 paralogs in *exo70A1* mutant background. **C-E** - EXO70 paralogs from Klebsormidium and Marchantia are not able to complement other prominent phenotypes previously found in EXO70.2 subfamily. Trichome *exo70H4* callose deposition defect (**C,D**) and *exo70B1* anthocyanin accumulation defect (**E**) could not be restored.

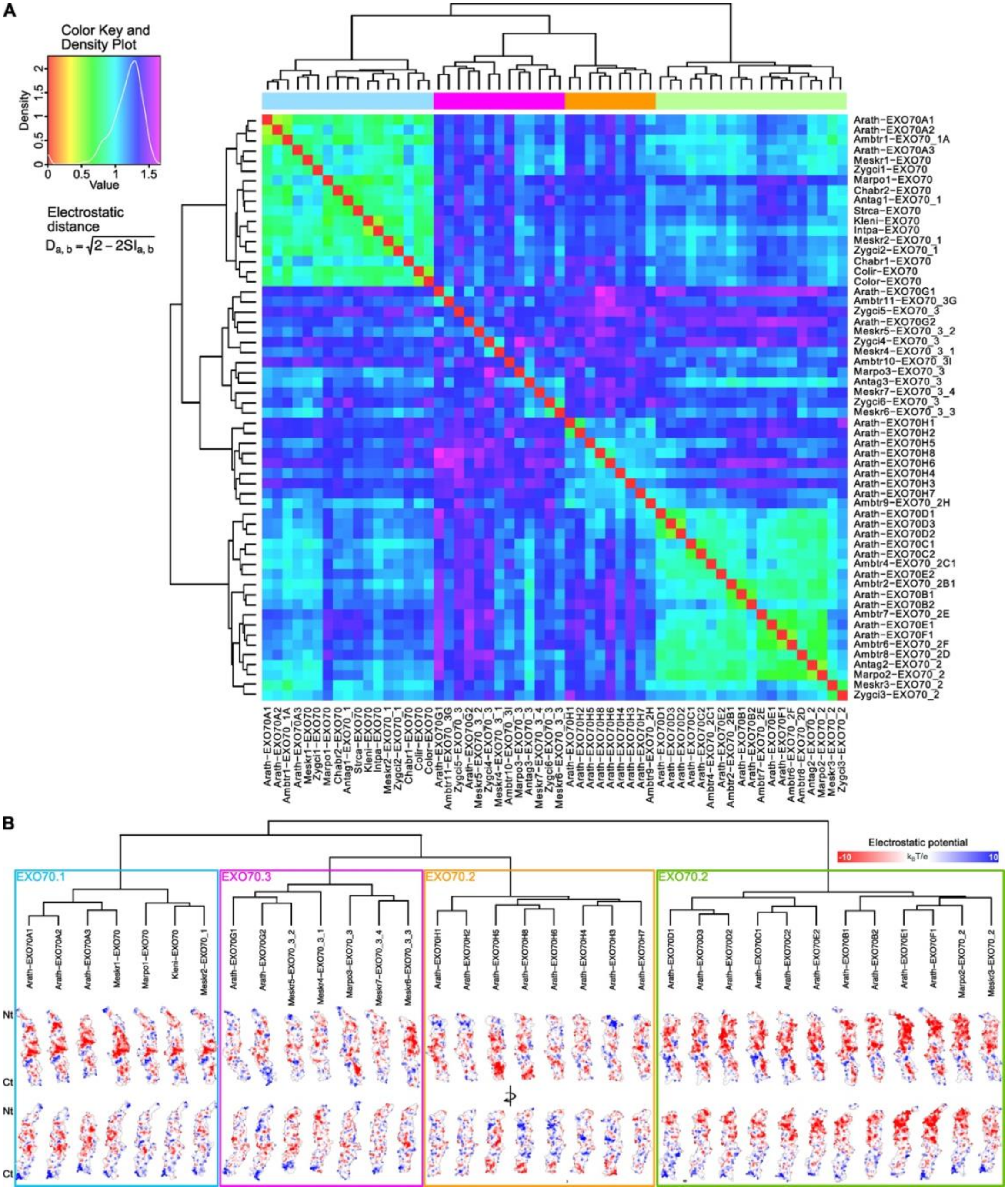


**Fig. S5. Electrostatic potential clustering of EXO70 paralogs across streptophytes**. **A** - Pairwise electrostatic similarity matrix of EXO70 proteins calculated using PIPSA (Protein Interaction Property Similarity Analysis; https://pipsa.h-its.org/pipsa/). Electrostatic distances (D) were computed from Poisson–Boltzmann–derived surface potentials and clustered hierarchically. Warmer colors indicate higher electrostatic similarity, whereas cooler colors indicate greater divergence. Clustering reveals that EXO70 proteins group primarily according to electrostatic surface properties rather than strict phylogenetic proximity, with canonical EXO70.1-like proteins from streptophyte algae and land plants clustering together despite substantial sequence divergence. **B** - Representative electrostatic surface potential maps of EXO70 proteins corresponding to major clusters identified in panel (A). Electrostatic potentials are displayed on the molecular surface (–10 to +10 kBT/e), oriented consistently from N-terminus (Nt) to C-terminus (Ct). Colored boxes denote clusters with shared electrostatic signatures, illustrating conservation of surface charge patterns in canonical EXO70.1-like proteins and pronounced divergence in EXO70.2- and EXO70.3-associated clusters.

Ambtr – *Amborella trichopoda,* Antag – *Anthoceros agrestis,* Arath – *Arabidopsis thaliana,* Chabr – *Chara braunii,* Chlat – *Chlorokybus atmophyticus,* Colir – *Coleochaete irregularis,* Color – *Coleochaete orbicularis,* Intpa – *Interfilum paradoxum,* Kleni – *Klebsormidium nitens,* Marpo – *Marchantia polymorpha,* Mesen – *Mesotaenium endlicherianum*, Mesvi – *Mesostigma viride,* Strca – *Streptofilum capillatum,* Zygci – *Zygnema circumcarinatum*

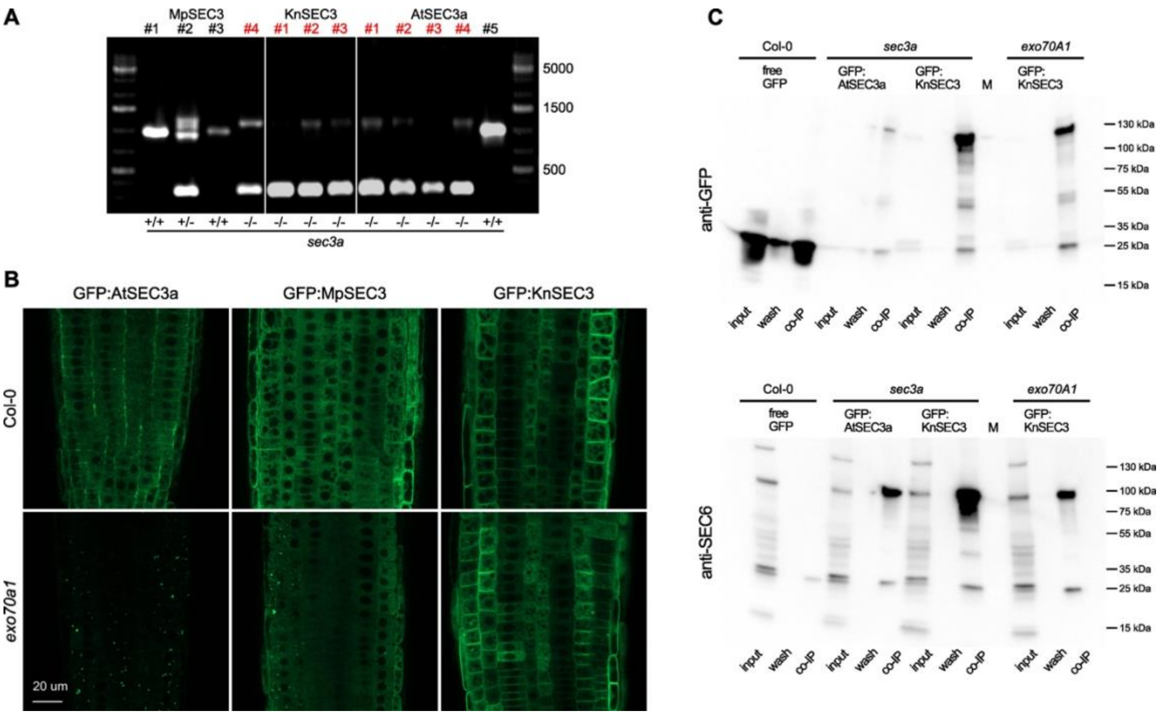


**Fig. S6. Complementation of *sec3a* mutant by SEC3 orthologs, localization of SEC3 orthologs in Arabidopsis wild-type and *exo70A1*mutant backgrounds and co-immunoprecipitation analysis. A** - Genotyping of progeny of gametophytically lethal *sec3a* +/- heterozygous mutant transformed by Arabidopsis SEC3a, Marchantia SEC3, and Klebsormidium SEC3 orthologs. Lines depicted in red shows successful complementants. **B** - Localization of strongly expressed GFP-tagged SEC3 isoforms from Arabidopsis, Marchantia and Klebsormidium in wild-type and *exo70A1* mutant background. Note the ectopic PM localization of KnSEC3 in both wild-type and *exo70A1* mutant background. **C** - Source image of the co-immunoprecipitation of the exocyst core subunit SEC6 with GFP-tagged AtSEC3a and KnSEC3 in wild-type, *sec3a*, and *exo70A1* mutant backgrounds shown in Fig 6C. Detection of SEC6 in SEC3 immunoprecipitates confirms association of both Arabidopsis and Klebsormidium SEC3 with the exocyst complex, including in the *exo70A1* background.

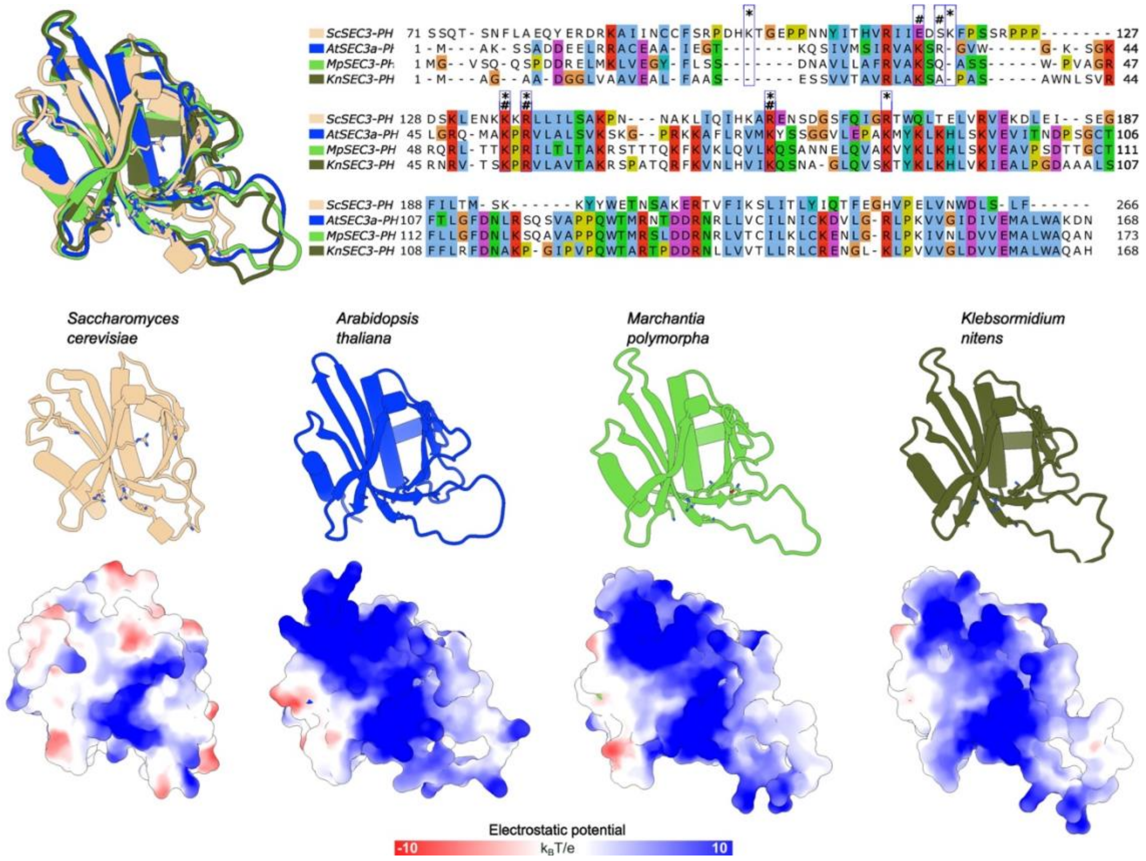


**Fig. S7. Structural analysis of PH domains from Arabidopsis, Marchantia and Klebsormidium SEC3 orthologs.** Top, Structural sequence alignment of PH domains from yeast, Arabidopsis, Marchantia and Klebsormidium SEC3 orthologs. Boxes depict amino acids involved in phosphoinositide binding in yeast (*) or Arabidopsis (#). Bottom, Structural and molecular surface models of selected SEC3-PH domains illustrating electrostatic potential distributions. Surface charge is displayed from negative (red) to positive (blue) electrostatic potential. Conserved residues forming phosphoinositide binding pocket are shown with side chains.

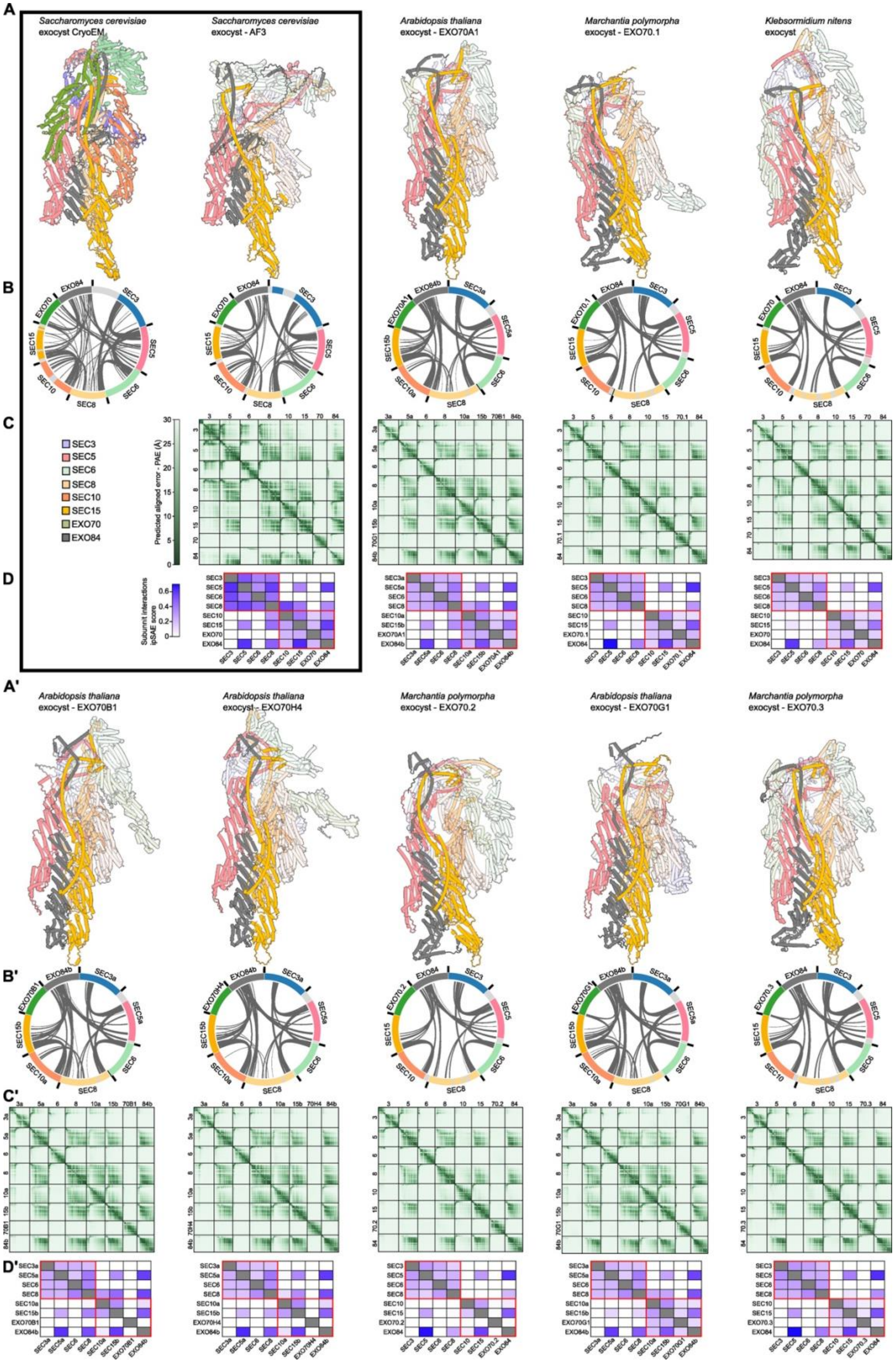
A
Saccharomyces cerevisiae exocyst CryoEM
Saccharomyces cerevisiae exocyst - AF3
Arabidopsis thaliana exocyst - EXO70A1
Marchantia polymorpha exocyst - EXO70.1
Klebsormidium nitens exocyst
B
C
D
SEC3
SEC5
SEC6
SEC8
SEC10
SEC15
EXO70
EXO84
Predicted aligned error - PAE (Å)
Subunit interactions ipSAE score
A'
Arabidopsis thaliana exocyst - EXO70B1
Arabidopsis thaliana exocyst - EXO70H4
Marchantia polymorpha exocyst - EXO70.2
Arabidopsis thaliana exocyst - EXO70G1
Marchantia polymorpha exocyst - EXO70.3
B'
C'
D'

**Figure S8. The validation of AF3 comparative modelling approach and *in silico* structural analysis of plant exocyst complex variants.** **A, A'** - schematic representations of the experimental CryoEM structure of partial exocyst from *Saccharomyces cerevisiae* and Alphafold3 (AF3)-based exocyst model variants from *S. cerevisiae*, *A. thaliana*, *M. polymorpha* and *K. nitens*. Subunits with uncertain global position are shown as semi-transparent. **B, B'** - chord plot projection of inter-subunit contacts. For experimental structure, all inter-subunit residue-residue contacts with distance < 5 Å are depicted, for AF3 models, inter-subunit residue-residue contacts with distance < 5 Å and predicted aligner error (PAE) < 10 Å are indicated. Regions not resolved in the CryoEM structure or trimmed before modelling are shown in grey. **C, C'** - Predicted aligned error (PAE) plots from AF3 predictions showing model confidence in relative residue positioning, with low (dark green) and high (light green) PAE values indicating robustly predicted rigid regions and versus flexible or uncertain regions, respectively. **D, D'** - Heatmaps showing the ipSAE (interaction prediction Score from Aligned Errors) for the individual subunit-subunit interfaces computed with the 10 Å threshold. Red rectangles indicate the interactions within module I and II, respectively.

Boxed panel - comparative analysis of the experimentally determined partial cryo-EM *S. cerevisiae* exocyst structure (Mei *et al*., 2018) with a near full-length model of the *S. cerevisiae* exocyst (6,407 of 7,334 amino acids) generated via template-free AF3 prediction. The AF3 model strongly supported a two-module organization of the complex, as indicated by the predicted aligned error (PAE) plots and the interaction prediction Score from Aligned Errors (ipSAE) metric (Dunbrack *et al*., 2025). Global alignment with USalign (Zhang *et al*., 2022) yielded a TM-score of 0.52 and an RMSD of 29 for the full complex, indicating overall agreement but notable differences for SEC3, SEC6, and EXO70, consistent with their PAE profiles. In contrast, PAE and ipSAE analyses identified SEC5, SEC8, SEC10, SEC15, and EXO84 as the structural core of the complex. Correspondingly, the TM-score for these core subunits increased to 0.66 and the RMSD decreased to 22, corroborating the robustness of their arrangement. Collectively, these results support the validity of template-free AF3 predictions for comparative structural analysis of exocyst complexes.

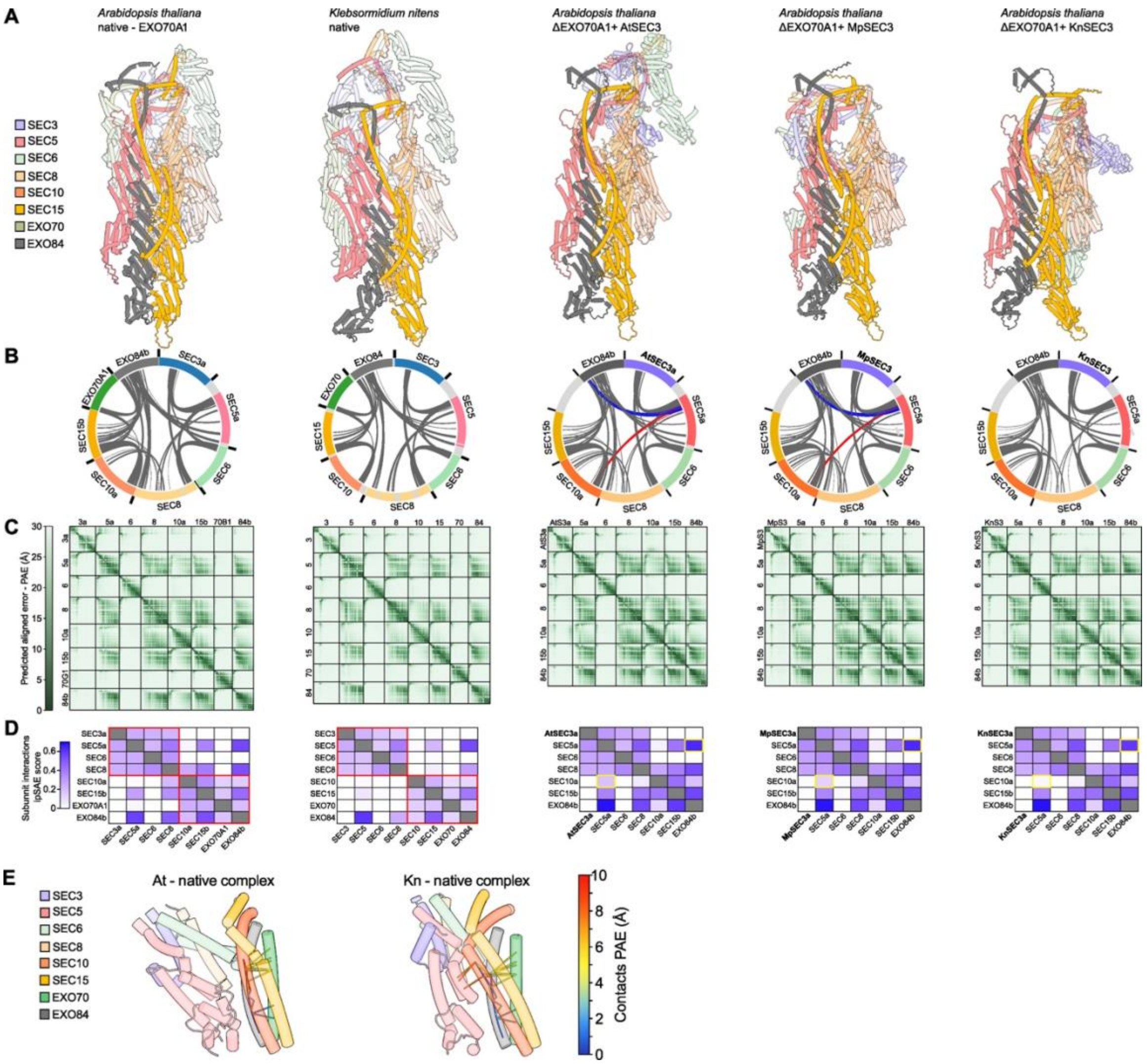

**Figure S9. Comparative *in silico* structural analysis of native exocyst models and models lacking EXO70A1 subunit.** **A** - schematic representations of the Alphafold3 (AF3)-based exocyst model native variants from *A. thaliana* and *K. nitens and* Arabidopsis variants lacking EXO70A1 and harboring SEC3 from Arabidopsis, Marchantia, or Klebsormidium. Subunits with uncertain global position are shown as semi-transparent. **B** - chord plot projection of inter-subunit contacts. Inter-subunit residue-residue contacts with distance < 5 Å and predicted aligner error (PAE) < 10 Å are indicated. Regions trimmed before modelling are shown in grey. **C** - Predicted aligned error (PAE) plots from AF3 predictions showing model confidence in relative residue positioning, with low (dark green) and high (light green) PAE values indicating robustly predicted rigid regions and versus flexible or uncertain regions, respectively. **D** - Heatmaps showing the ipSAE (interaction prediction Score from Aligned Errors) for the individual subunit-subunit interfaces computed with the 10 Å threshold. Red rectangles indicate the interactions within module I and II, respectively. **E** - Close-up visualization of selected subunit-subunit interactions in native Arabidopsis and Klebsormidium models. Note that for the sake of comparative clarity, part of panels A-D containing data for native Arabidopsis and Klebsormisium models are identical to those in Supplementary Fig. S8A-D.

**Table S1.** Database source

| Group | Abbrev. | Species | Type | Source | Reference |
|---|---|---|---|---|---|
| *Opisthokonts* | - | *Saccharomyces cerevisiae ATCC 204508/S288c* | Proteome | https://www.uniprot.org | - |
| | - | *Schizosaccharomyces pombe 972/ATCC 24843* | Proteome | https://www.uniprot.org | - |
| | - | *Homo sapiens* | Proteome | https://www.uniprot.org | - |
| | - | *Danio rerio* | Proteome | https://www.uniprot.org | - |
| | - | *Drosophila melanogaster* | Proteome | https://www.uniprot.org | - |
| Chlorophytic algae | - | *Chlamydomonas reinhardtii CC-4532* | Genome v5.6/v6.1 | https://phytozome.jgi.doe.gov | https://doi.org/10.1126/science.1143609 |
| | - | *Volvox carteri* | Genome v2.1 | https://phytozome.jgi.doe.gov | https://doi.org/10.1126/science.1188800 |
| | - | *Chlorella sorokiniana* | TSA | NCBI BioProject: PRJNA218510 | Li, unpubl. |
| | - | *Chlorella sp. HS2* | TSA | NCBI BioProject: PRJNA590327 | Pierrelee, unpubl. |
| | - | *Chlorella variabilis* | mRNA | NCBI provisional RefSeq | https://doi.org/10.1105/tpc.110.076406 |
| | - | *Coccomyxa subellipsoidea C-169* | Genome v2.0 | https://phytozome.jgi.doe.gov | https://doi.org/10.1186/gb-2012-13-5-r39 |
| | - | *Chromochloris zofingiensis* | Genome v5.2.3.2 | https://phytozome.jgi.doe.gov | https://doi.org/10.1073/pnas.1619928114 |
| *Mesostimatophyceae* | Chlat | *Chlorokybus atmophyticus CCAC 0220*<br>*Chlorokybus atmophyticus* | Genome<br>TSA | http://ftp.cngb.org/<br>OneKP SampleID: AZZW | https://doi.org/10.1038/s41477-019-0560-3<br>https://doi.org/10.1038/s41586-019-1693-2 |
| | Mesvi | *Mesostigma viride CCAC 1140*<br>*Mesostigma viride NIES-296*<br>*Mesostigma viride NIES-995* | Genome<br>Genome<br>TSA | http://ftp.cngb.org/<br>NCBI BioProject: PRJNA505414<br>NCBI BioProject: PRJNA242253 | https://doi.org/10.1038/s41477-019-0560-3<br>https://doi.org/10.1002/advs.201901850<br>https://doi.org/10.1038/nplants.2014.4 |
| | Spimi | *Spirotaenia minuta* | TSA | OneKP SampleID: NNHQ | https://doi.org/10.1038/s41586-019-1693-2 |
| *Klebsormidiophyceae* | Strca | *Streptofilum capillatum* | TSA | https://figshare.com/s/1b1a1672c26a96f10ee8 | https://doi.org/10.1101/2024.03.08.584070 |
| | Kleni, Kn | *Klebsormidium nitens NIES-2285* | Genome | http://www.plantmorphogenesis.bio.titech.ac.jp | https://doi.org/10.1038/ncomms4978 |
| | Intpa | *Interfilum paradoxum* | TSA | OneKP SampleID: FPCO | https://doi.org/10.1038/s41586-019-1693-2 |
| *Charophyceae* | Chabr | *Chara braunii S276* | Genome<br>TSA | https://bioinformatics.psb.ugent.be/<br>NCBI BioProject: PRJNA492241 | https://doi.org/10.1016/j.cell.2018.06.033<br>https://doi.org/10.1111/nph.15829 |
| | Nitmi | *Nitella mirabilis* | TSA | NCBI BioProject: PRJNA242253, PRJNA158153 | Ju *et al.*, unpubl.; Thierer *et al.*, unpubl. |
| *Coleochaetophyceae* | Color | *Coleochaete orbicularis* | TSA | NCBI BioProject: PRJNA242253 | Ju *et al.*, unpubl. |
| | Colir | *Coleochaete irregularis* | TSA | OneKP SampleID: QPDY | https://doi.org/10.1038/s41586-019-1693-2 |
| *Zygnematophyceae* | Mesen | *Mesotaenium endlicherianum SAG 12.97* | Genome<br>TSA | https://figshare.com/<br>OneKP SampleID: WDCW | https://doi.org/10.1016/j.cell.2019.10.019<br>https://doi.org/10.1038/s41586-019-1693-2 |
| | Mesbr | *Mesotaenium braunii* | TSA | OneKP SampleID: WSJO | https://doi.org/10.1038/s41586-019-1693-2 |
| | Meste | *Mesotaenium testaceovaginatum* | TSA | NCBI BioProject: PRJEB72628 | Busch, unpubl. |
| | Meskr | *Mesotaenium kramstae NIES-658* | Genome v2.0<br>TSA | https://phycocosm.jgi.doe.gov<br>OneKP SampleID: NBYP | https://doi.org/10.2216/i0031-8884-4-1-23.1<br>https://doi.org/10.1038/s41586-019-1693-2 |
| | Zygci | *Zygnema circumcarinatum SAG698-1b*<br>*Zygnema circumcarinatum SAG698-1a* | Genome<br>Genome | https://bcb.unl.edu/Zygnema_4genomes/SAG698-1b/<br>https://bcb.unl.edu/Zygnema_4genomes/SAG698-1a/ | https://doi.org/10.1038/s41588-024-01737-3<br>https://doi.org/10.1038/s41588-024-01737-3 |
| | Cylbr | *Cylindrocystis brebissonii* | TSA | OneKP SampleID: YOXI, VAZE | https://doi.org/10.1038/s41586-019-1693-2 |
| Bryophyta | Antag | *Anthoceros agrestis* | TSA | OneKP SampleID: TWUW, BSNI | https://doi.org/10.1038/s41586-019-1693-2 |
| | Leidu | *Leiosporoceros dussii* | TSA | OneKP SampleID: ANON | https://doi.org/10.1038/s41586-019-1693-2 |
| | Porna | *Porella navicularis* | TSA | OneKP SampleID: KRUQ | https://doi.org/10.1038/s41586-019-1693-2 |
| | Marpo, Mp | *Marchantia polymorpha* | Genome v3.1 | https://phytozome.jgi.doe.gov | https://doi.org/10.1016/j.cell.2017.09.030 |
| | Sphfa | *Sphagnum fallax* | Genome v0.5 | https://phytozome.jgi.doe.gov | DOE Joint Genome Institute |
| | Phypa | *Physcomitrium patens* | Genome v3.3 | https://phytozome.jgi.doe.gov | https://doi.org/10.1111/tpj.13801 |
| Lycopodiophyta | Selmo | *Selaginella moellendorffii* | Genome v2.1 | https://phytozome.jgi.doe.gov | https://doi.org/10.1126/science.1203810 |
| | Selse | *Selaginella sellowii* | TSA | NCBI BioProject: PRJNA625275 | Alejo-Jacuinde *et al.*, unpubl. |
| | Isote | *Isoetes tegetiformans* | TSA | OneKP SampleID: PKOX | https://doi.org/10.1038/s41586-019-1693-2 |
| | Isota | *Isoetes taiwanensis* | Genome | https://genomevolution.org/ | https://doi.org/10.1038/s41467-021-26644-7 |
| | Lyccl | *Lycopodium clavatum* | Genome | GWH Accession No. GWHBJYW00000000; https://figshare.com/ | https://doi.org/10.1007/s11103-023-01366-0 |
| | Dipco | *Diphasiastrum complanatum* | Genome v3.1 | https://phytozome.jgi.doe.gov | DOE Joint Genome Institute |
| Monilophyta | Cerri | *Ceratopteris richardii* | Genome v2.1<br>TSA | https://phytozome.jgi.doe.gov<br>NCBI BioProject: PRJNA511033 | https://doi.org/10.1038/s41477-022-01226-7<br>https://doi.org/10.1038/s41598-019-53968-8 |
| | Ptevi | *Pteris vittata* | TSA | NCBI BioProject: PRJNA492307 | Cai *et al.*, unpubl. |
| Gymnosperms | Ginbi | *Ginkgo biloba* | Genome<br>TSA | http://gigadb.org/dataset/100613<br>NCBI BioProject: PRJNA270069, PRJNA515544 | https://doi.org/10.5524/100613<br>Han *et al.*, unpubl.; Zou *et al.*, unpubl. |
| | Thupl | *Thuja plicata v3.1* | Genome v3.1 | https://phytozome.jgi.doe.gov | https://doi.org/10.1101/gr.276358.121 |
| Angiosperms | Ambtr | *Amborella trichopoda* | Genome v1.0<br>Genome v2.1<br>TSA | https://phytozome.jgi.doe.gov<br>https://phytozome.jgi.doe.gov<br>OneKP SampleID: EQDA | https://doi.org/10.1126/science.1241089<br>The Open Green Genomes Initiative,<br>https://doi.org/10.1038/s41586-019-1693-2 |
| | Bradi | *Brachypodium distachyon* | Genome v3.2 | https://phytozome.jgi.doe.gov | https://doi.org/10.1038/nature08747 |
| | Arath, At | *Arabidopsis thaliana* | Genome Araport11 | https://phytozome.jgi.doe.gov | https://doi.org/10.1111/tpj.13415 |

**Table S2.** List of PCR primers used in this study

| ID | Primer name | Primer sequence 5'-3' |
|---|---|---|
| P1 | AOC.GWB101_F | GATCCCCCGGGATAGGTTTAATTAAGAGCT |
| P2 | AOC.GWB101_R | CTTAATTAAACCTATCCCGGGG |
| P3 | HindIII.*proMpEF1*_F | AAGCTTCCGACAAGCAGAAAAGGAAACTT |
| P4 | NcoI.*proMpEF1*_R | CCATGGTGACAACCTTTCTGCAGGC |
| P5 | HindIII.*proAtUBQ10*_F | TATAAGCTTGTCGACGAGTCAGTAATAAACG |
| P6 | CTRQ_R | GCTGAACTTGTGGCCGTTTA |
| P7 | NcoI.GFP_F | CCATGGTGAGCAAGGGCGAGGAGCTGTT |
| P8 | BamHI.GFP_R | GGATCCGCCACCAGATCCACCCTTGTACAGCTCGTCCATGCC |
| P9 | BglII.KnSEC3_F | AGATCTGGAGGTATGGCTGGCGCTGCGGAC |
| P10 | PacI.KnSEC3_R | TTAATTAATCACACTGTTCGGAACAGTTCGCGCATC |
| P11 | BamHI.KnSEC15_F | TAGGGATCCGGAGGTATGGCGGTCAACAGGGG |
| P12 | PacI.KnSEC15_R | ATTTAATTAACGGTTGCACGGGTGTTATCT |
| P13 | BglII.KnEXO70_F | AGATCTGGAGGTATGGGCGCCATGGAGGTAGAACGTCTGGTC |
| P14 | PacI.KnEXO70_R | TTAATTAATTACCTTCGGTTTCCTCCCTCAA |
| P15 | XmaI.MpSEC3_F | CCCGGGGGAATGGGGGTTTCTCAGCAGAGTCC |
| P16 | PacI.MpSEC3_R | TTAATTAACTATACATTCTTTAAAAGCTCCTTCATTTCGGCAGAC |
| P17 | BamHI.MpSEC15_F | GGATCCGGAGGTATGATGAACGCCAAAGGAGGG |
| P18 | PacI.MpSEC15_R | TTAATTAATTACAACTGATCTCTCAATCTTCTACACAGGG |
| P19 | BglII.MpEXO70.1_F | AGATCTGGAGGTATGGGTGCGGCGGCAGACATAGAG |
| P20 | PacI.MpEXO70.1_R | TTAATTAATTACCGTCTTTGATCAGCTCGCAT |
| P21 | BamHI.MpEXO70.2_F | GGATCCGGAGGTATGGCCGCATTTGATGGGGAAGA |
| P22 | PacI.MpEXO70.2_R | TTAATTAATCAGCTGGAGGAAAAAGATTTCCTTCTCAG |
| P23 | BglII.MpEXO70.3_F | AGATCTGGAGGTATGGGGCTGGATAGCTCGG |
| P24 | PacI.MpEXO70.3_R | TTAATTAATCATCCGTCCCCAAGGTTGCCGTCAGTCGTTCCGTTGCGCGGTTTACT |
| P25 | AgeI.AtSEC3A_F | TAAACCGGTATGGCGAAATCAAGCGCC |
| P26 | PacI.AtSEC3A_R | GCTTTAATTAATTACATGGAAGCCAGAAGTCCTCT |
| P27 | BamHI.AtSEC15B_F | TAGGGATCCGGAGGTATGCAATCGTCGAAAGGACG |
| P28 | PacI.AtSEC15B_R | GCTTTAATTAATCAGCTCACATCTTTCAATCTCTTTA |
| P29 | XmaI.AtEXO70A1_F | ATCCCCGGGATGGCTGTTGATAGCAGAATGG |
| P30 | PvuI.AtEXO70A1_R | TACCGATCGTTACCGGCGTGGTTCATTCAT |
| P31 | exo70A1-1-LP | TGTCACCCACCAAACCAAACC |
| P32 | exo70A1-1-RP | TCCACAATAGCTTTGTTCCCTGAA |
| P33 | LB-new | GAACAACACTCAACCCTATCTCGGGC |
| P34 | sec3A-1-ins | TCTTATTCTCTCCCTCTCTTGGTTG |
| P35 | sec3A-1-wt | AACTGCCTTCAGTTATGGTTCTATACTT |
| P36 | GABI-T-DNA | CCCATTTGGACGTGAATGTAGACAC |
| P34 | exo70H4-LP | TGGGATTTCGTTTCACAGTTC |
| P35 | exo70H4-RP | AATCCGTCATGACGTTGGTAG |
| P36 | exo70B1-LP | CGTGGCAGGAGTTAGAAGATG |
| P37 | exo70B1-RP | TTGTCTGCGTTTTTCCCTATG |
| P38 | sec15B-LP | AAGCATCCGTTTCAGCGTTGAC |
| P39 | sec15B-RP | GTTGAATTCCTATGGTTTCTAGAAAAC |

### Legends for Datasets

#### Dataset S1-S2 (separate files).

**Dataset S1 (separate file).** Deposited full EXO70 phylogenetic tree (EXO70_tree) presented in Figure 1. along with the matrix used for calculations (EXO70_matrix). Folder also contains alternative EXO84 tree (EXO84_tree) again with matrix used for this calculation (EXO84_matrix) – supports of which are partially transferred into Supplementary Figure S1I in form of numbers in brackets (note, that it`s without Setaphyta (liverworts and mosses) and Opisthokonta/Chlorophyta as an outgroup).

**Dataset S2 (separate file).** Datasets of full-length protein sequences for 8 exocyst complex subunits used in this study, along with matrices used for the phylogenetic tree calculations presented in Supplementary Figure S1A-I.
